# Evolutionary Principles of Bacterial Signaling Capacity and Complexity

**DOI:** 10.1101/2022.03.11.484041

**Authors:** Ran Mo, Yugeng Liu, Yuanyuan Chen, Yingjin Mao, Beile Gao

## Abstract

Microbes rely on signal transduction systems to sense and respond to environmental changes for survival and reproduction. It is generally known that niche adaptation plays an important role in shaping the signaling repertoire. However, the evolution of bacterial signaling capacity lacks systematic studies with a temporal direction. Particularly, it is unclear how complexity evolved from simplicity or vice versa for signaling networks. Here we examine the evolutionary processes of major signal transduction systems in *Campylobacterota* (formerly *Epsilonproteobacteria*), a phylum with sufficient evolutionary depth and ecological diversity. Evolution of signaling systems within *Campylobacterota* shows two opposite trends. During niche expansion, signaling complexity increases with gene expansions through horizontal gene transfer (HGT), gene duplication, fusion and fission, which create opportunities for genetic innovation and pathway integration. In contrast, as the lineages adapt to a specialized niche, complexity decreases with massive gene losses that lead to the decline or disappearance of pathways mediated by multiple transmitters. Overall, signaling capacity and complexity arise and drop together in *Campylobacterota*, determined by sensory demand, genetic resources and co-evolution within the genomic context. These findings reflects plausible evolutionary principles for other cellular networks and genome evolution of the *Bacteria* domain.

## INTRODUCTION

Five decades of vigorous studies have elucidated major signal transduction systems in bacteria and they came into the limelight in the following order:

1. Chemosensory system (also called the chemotaxis system), mostly involved in navigation of motility ^1^;
2. Two-component system (TCS), primarily for regulation of gene expression ^2,3^;
3. Secondary messenger dependent systems represented by cyclic adenosine monophosphate (cAMP) ^4^, cyclic diguanylate (c-di-GMP) ^5^, cyclic diadenylate (c-di-AMP) ^6^ and more ^7,8^;
4. Serine/Threonine/Tyrosine protein kinases (STYKs) and protein phosphatases mediated pathways ^9^;
5. Extracytoplasmic function σ factors (ECFs) as alternative σ factors in redirection of transcription ^10^;
6. Quorum sensing (QS), in cell to cell communication ^11^.

There are also one-component systems (OCSs), each composed of a single protein that works by domain-domain interaction, but OCS proteins often become components of other systems once linked with other proteins ^12^. While each signal transduction system may stand on its own, they are also intertwined to form a network that confers fitness advantage for bacterial adaptation to the ever-changing environment. Previous census studies have counted and sorted the components in each system across diverse bacterial lineages, learning their system composition and diversity, modularity and plasticity, phylogenetic distribution, correlation with genome size and ecology, and so on ^13-19^. However, beyond the single system level, how the signaling capacity of bacterial species evolved remains largely elusive. This general question can be specified as: 1. How are new components and network links added to increase system complexity? 2. What kind of genetic innovations accompany the increase in the number of components? 3. Why are some systems abundant but some are absent in different bacterial species? Answers to these questions are fundamental to understand the organization of any biological network, and also to shed light on the evolution of genome that encodes a supernet of all cellular processes.

A systematic approach to tackle the above questions would be to track the evolution of the major signal transduction systems in the *Bacteria* based on the species/lineage branching order. Analyses of a large number of diverse bacterial species with finely spaced phylogenetic progression are needed for a full coverage of signaling capacity at different complexity levels. In addition, it is believed that ecological adaptation is the driving force for signaling network evolution, thus species habitat should be considered when comparing the dynamics of the signaling proteins in different species ^20^. Currently, it is not realistic to analyze the entire *Bacteria* domain with the enormous number of sequenced genomes, simply because there is no consensus on the root of the bacterial tree and the order of phyla divergence ^21,22^. In addition, since the above questions focus on the evolutionary processes rather than the origin of signal transduction systems, a starting point is needed but not necessarily the one tracing back to the last bacterial common ancestor. Instead, a monophyletic group with a deep root, well-supported phylogeny and wide ecological distribution would provide a good representation. We propose that *Campylobacterota* (phyl. nov.), previously *Epsilonproteobacteria* class, is a good model for such a pilot study ^23^.

## RESULTS

### An eco-evo framework from deep-sea to human gut

*Campylobacterota* is one of the phyla with many complete genomes of diverse species based on current NCBI and GTDB taxonomy and genome collections ^24,25^. Other bacterial phyla with more than 50 completely sequenced species are either not monophyletic or lack ecological diversity, thus not selected for this study (see details in Supplementary Table 1). Both 16S rRNA trees and phylogenomic analyses strongly suggest that the ancestral lineage of this phylum was composed of specialist organisms residing at the deep-sea hydrothermal vents, which are a model system for studying the Archaean Earth environment ^23,26^. We reconstructed the phylogeny for all species with completely sequenced genomes from *Campylobacterota* and collected their ecological and physiological details (Fig. 1a, Extended Data Fig. 1 and Supplementary Table 2). The deepest branch of this phylum consists of strictly anaerobic and thermophilic chemolithoautotrophs that were isolated only from the deep-sea hydrothermal vents. Their metabolic characteristics and prevalence in the vent microbial community suggest that the ancient *Campylobacterota* may have played an important role in the Early Earth history ^26^. Then, later lineages underwent niche expansion towards the surface of the ocean and terrestrial water reservoirs; these lineages can be grouped as free-living generalists. Along with ecological diversification, they display physiological transitions from anaerobic to microaerobic/aerobic environment, thermophily to mesophily, and autotrophy to heterotrophy. Furthermore, some lineages became host-associated commensals or pathogens represented by two genera, *Campylobacter* and *Helicobacter*, obligately mesophilic and heterotrophic. Eventually, some species became restricted to specific hosts such as *Helicobacter pylori*, successful colonizers of human gut.

**Fig. 1.**
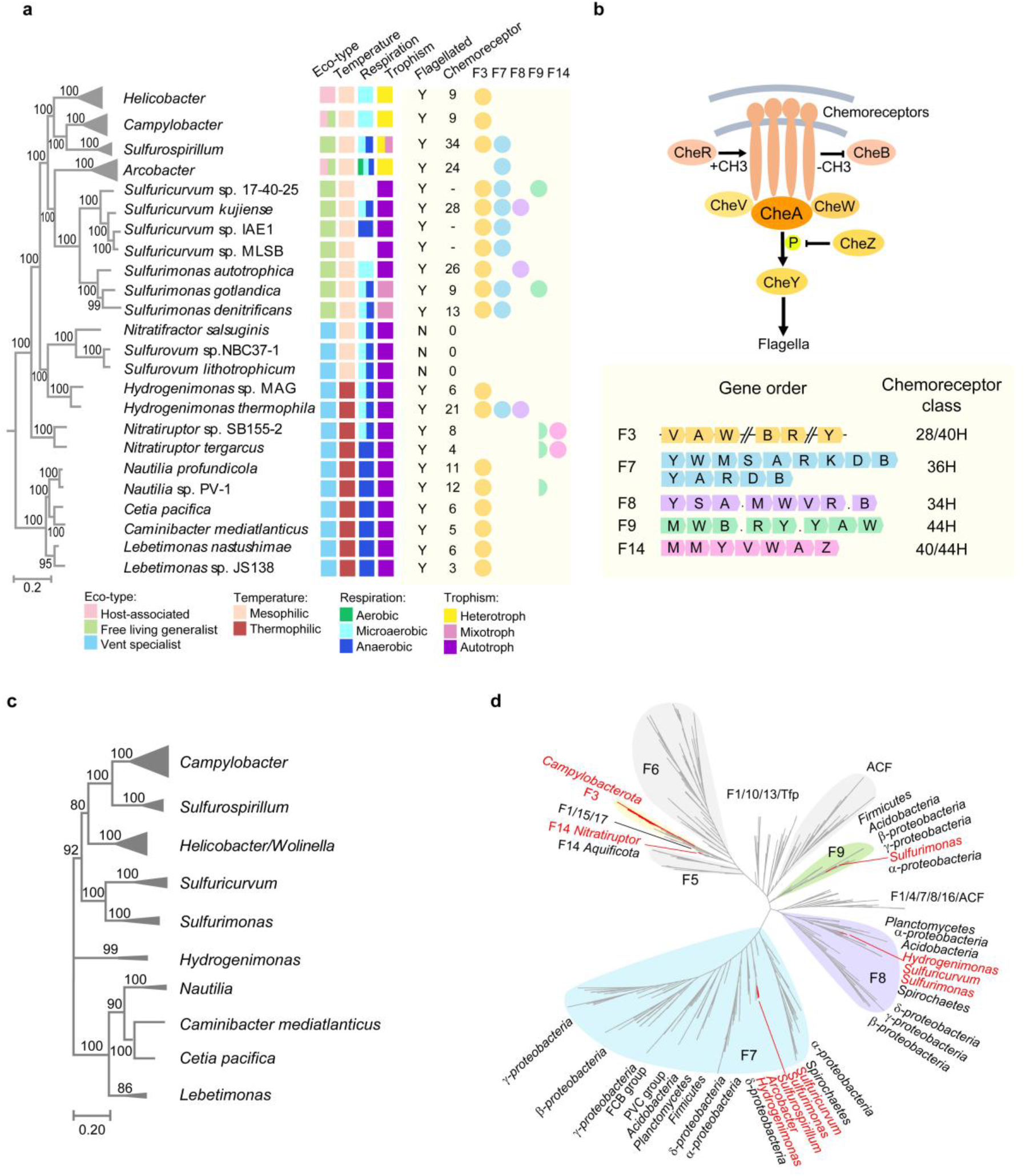
Chemosensory system in *Campylobacterota*. **(*a*)** Phylogeny (left), ecophysiological characteristics (middle), chemosensory system (right) of the *Campylobacterota*. Tree topology is based on Extended Data Fig. 1 and collapsed for 4 genera with multiple sequenced species but the same chemosensory F class pattern. “-” in the chemoreceptor column indicates that the total number of chemoreceptor genes is unavailable due to genome incompleteness. Full circle on the right represents that the full gene set of specific F class is present; half circle means only half components of the F class are present. **(*b*)** Model of chemosensory system and characteristics of F3/F7/F8/F9/F14 classes in terms of composition, gene order, and chemoreceptor type. Abbreviations in gene order: A, *cheA*; B, *cheB*; D, *cheD*; K, a gene encoding histidine kinase; M, chemoreceptor; R, *cheR*; S, a gene encoding STAS domain alone; V, *cheV*; W, *cheW*; Y, *cheY*; “.” represents hypothetical proteins. **(c)** Phylogeny of F3 class core proteins (CheA, CheV and CheW) in *Campylobacterota*. **(*d*)** Phylogeny of CheA homologs of distinct F classes from diverse bacterial phyla. The *Campylobacterota* phylum and its genera are highlighted in red.

It is clear that *Campylobacterota* has a deep evolutionary origin and a traceable niche expansion from deep-sea hydrothermal vents to diverse habitats of modern lineages, making this phylum suitable for inferring evolutionary processes by assessment of its contemporary diversity. Moreover, species of this phylum have a narrow range of genome size variation (1.4∼3.5 Mb), without dramatic genome reduction or expansion largely due to random genetic drift (Supplementary Table 2). This enables examination of ecological adaptive selection on genome evolution in a stepwise manner. Here, based on this “eco-evo” framework, we dissect each signal transduction system by following their compositional changes, compare their evolutionary processes and identify their determinants in the signaling network.

### Chemosensory system: add, reduce but rarely stir

The chemosensory system is composed of multicomponent pathways that sense environmental stimuli by different chemoreceptors and integrate all the signals to affect the activity of CheA kinase ^27^. Activated CheA can autophosphorylate and pass on the phosphoryl group to its response regulator CheY to control motility, biofilm formation, development or other cellular processes (Fig. 1b). The ultimate complexity of this system is reflected by the presence of several chemosensory pathways/arrays with different composition and structure in some bacterial cells ^28,29^. A milestone classification scheme for chemosensory system has sorted them into 19 classes based on gene order, domain architecture, auxiliary component(s), and chemoreceptor types ^30^. These classes include one type IV pili class (Tfp), one “alternative cellular functions” class (ACF), and 17 flagellar classes (F1-F17) ^30^. We followed this classification scheme to analyze all chemosensory genes in *Campylobacterota* species and identified 5 classes (F3/F7/F8/F9/F14) in this phylum. Their distinctive gene order/components within chemosensory gene clusters and corresponding chemoreceptor types are summarized in Fig. 1b.

Mapping all identified chemosensory classes in the “eco-evo” framework of *Campylobacterota*, we found that F3 class is conserved throughout *Campylobacterota* from hydrothermal vent specialists to host-associated species, except those that are either unflagellated or have distinctive flagellar gene sets (Fig. 1a, Extended Fig. 2 and unpublished paper by Mo et al). The F3 class is defined by the presence of a core gene operon *cheVAW*, additional receiver (REC) domain in CheA kinase, and an auxiliary CheB lacking the REC domain ^30^ (Fig. 1b). A phylogenetic tree based on core proteins of F3 class shows the same branch pattern as the species tree, suggesting that F3 class evolved in the common ancestor of *Campylobacterota* and was vertically passed on to its descendants (Fig. 1c).

In addition to F3, other chemosensory classes are found in free-living generalists including F7, F8, or F9, coupled with a larger number of chemoreceptor genes (Fig. 1a). These additional F classes likely have been acquired by HGT since they branch with other bacterial phyla with the same F class in phylogenetic tree analysis (Fig. 1d). Consistently, these classes have all or most of their components encoded in a single cluster including their cognate chemoreceptors, facilitating genetic transfers within this phylum and from other bacterial phyla of the same habitat (Extended Fig. 2). Moreover, the narrow or sporadic species distribution patterns of these non-F3 classes suggest that they were introduced at different branching points of *Campylobacterota* lineages and subsequently lost in some species or transferred again within this phylum (Fig. 1a, d). Thus, in contrast to the vertically inherited F3 class, they are easily gained and lost by *Campylobacterota* during niche expansion, and completely absent in recently derived lineages associated with animal or human hosts.

To increase the sensing ability for niche expansion, bacteria could simply gain more chemoreceptors by gene duplication/divergence and HGT but using the same transmitter gene sets. However, most free-living generalists examined here acquired extra chemosensory classes, adding complexity to signal transmission while expanding sensory repertoire. Later, when the lineages became niche specialized, they only retain one F class with a reduced number of chemoreceptors, or lose the entire chemosensory system if their flagellar genes are lost. For example, species of the *Sulfurovum* genus that are either ubiquitous in deep-sea vent biofilm, or living as epibiotic symbionts of deep-sea animals, do not encode any chemosensory or flagellar genes, consistent with their sessile lifestyle (Fig. 1a) ^31^. These results suggest that the presence and complexity of chemosensory system is likely to be determined by sensory demand and signal output such as flagella. However, an “F” chemosensory class does not necessarily control flagellar motility and many signal outputs for species with multiple chemosensory classes are currently unknown ^28^. Future functional characterization of all signal outputs in species with complex chemosensory system will reveal specific determinants for each chemosensory class.

The above phylogenetic analyses clearly show that all the non-F3 classes in *Campylobacterota* are obtained by HGT, and not by duplication and divergence of the ancestral F3 class. Moreover, although different chemosensory classes within the same genome undergo dynamic gain and loss, their main transmitter genes rarely mix and match. In all the chemosensory classes of *Campylobacterota*, we only see two examples of mixed F class genes: F3-*cheZY* incorporated into the new F7 class in *Arcobacter*; and half F9 class without its own kinase that must function with the only F class in the species (unpublished paper by Mo et al). The fact that only 19 chemosensory classes could be identified among the great diversity of the *Bacteria* domain also indicates that genetic innovation of new chemosensory class is rare during bacterial evolution and species with multiple chemosensory class mainly gain extra transmitter sets by HGT. Thus, we conclude that the chemosensory system follows “add, reduce but rarely stir” evolutionary mode, compared to a description of genome evolution of bacterial pathogens - “add, stir and reduce” ^32^. Importantly, the intrinsic “rarely stir” feature ensures a traceable history, making the F class classification scheme work.

### TCS perplexes with more and more REC domains

The TCS is typically composed of a sensory histidine kinase (HK) and a response regulator (RR) ^33^. An HK protein generally has one or more sensor domain(s) at the N-terminal and HisKA plus HATPase domains at the C-terminal that can autophosphorylate upon activation by the sensor domain and then transfer the phosphoryl group to the REC domain of its cognate RR (Fig. 2a). These HK-RR pairs can hardly make a sophisticated network by increasing their numbers, in fact a few atypical HKs truly expand the diversity and complexity of TCS ^34^. Based on domain architecture, atypical HKs are classified into two types. One type is hybrid histidine kinase (HHK) with C-terminal REC domain that can perform a phosphorelay or tune its own kinase activity. Another type is hybrid response regulator (HRR) with N-terminal REC domain fused to transmitter domains HisKA and HATPase, and importantly the N-terminal REC domain is phosphorylated by another HK protein rather than its own kinase domain (Fig. 2a) ^35,36^. Because of the built-in REC domain, both HHKs and HRRs can participate in multicomponent or multistep signal transduction processes, but it is difficult to delineate their information flow in contrast to the paired HK-RR paradigm ^37,38^.

**Fig. 2.**
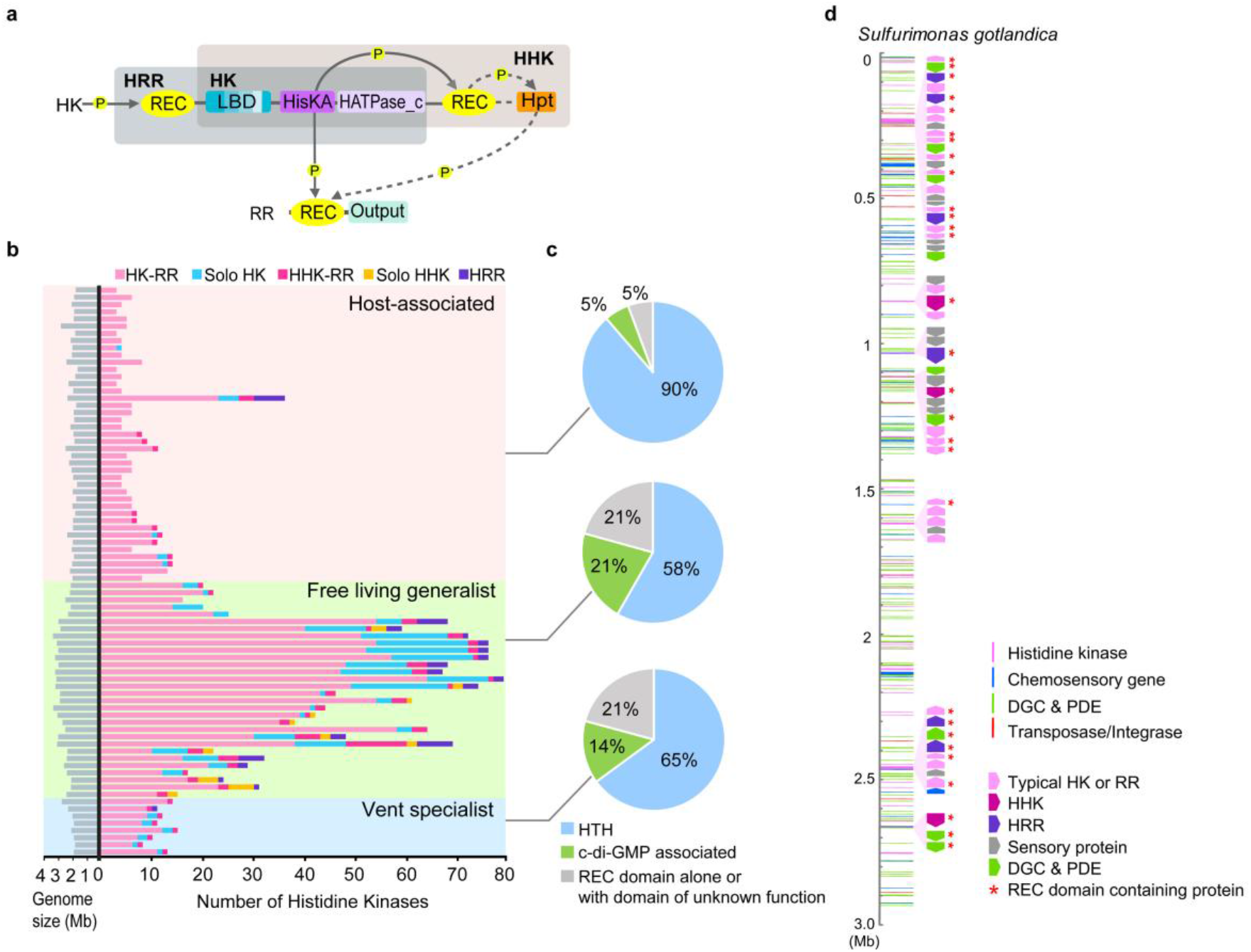
TCSs in *Campylobacterota*. **(*a*)** Schematic modular organization of TCS proteins. Phosphotransfer is depicted by solid arrow line and phosphorelay by dashed arrow line. Abbreviations for domains: REC: receiver domain; LBD: ligand-binding domain; HisKA: histidine kinase phosphoacceptor domain; HATPase_c: histidine kinase-like ATPase domain; Hpt: histidine-containing phosphotransfer domain. **(*b*)** The abundance of HKs in *Campylobacterota* species. The number details in Supplementary Table 3. **(*c*)** The proportions of output types of RRs in vent specialists, generalists and host-associated species of *Campylobacterota*. **(*d*)** Atypical HK enriched gene clusters in the genome of *Sulfurimonas gotlandica* GD1, which is linearized for display purpose. All signal transduction genes are represented as colored stripes based on their starting location and atypical HK enriched regions are labeled beside the genome.

To evaluate the abundance of TCS in *Campylobacterota* genomes, we counted all HK proteins with both HisKA and HATPase domains and found that the numbers from each species roughly correlate with their ecological breadth (Fig. 2b). Both vent specialists and host-associated groups have relatively fewer and typically classical TCS pairs (∼10 or less), while free-living generalists have fair amounts of solo HKs, HHKs and HRRs in addition to the paired HKs, with the peak number of 79 HKs in total. Accordingly, the number of RRs increases with that of HKs, and their output domains also diversify. Besides the RRs with the DNA-binding helix-turn-helix domain as the common output to regulate gene expression, 21% of RRs in free-living *Campylobacterota* generalists have c-di-GMP turnover enzymes as signal outputs and another 21% have other domains mostly of unknown function (Fig. 2c). As previously reported, TCS can easily integrate with other signaling systems through the fusion of REC domain with non-canonical modules ^36,39^.

These analyses highlighted two notable trends for HHKs and HRRs: their amount is positively related to the total number of HKs, and the genes encoding them are frequently flanked by TCS gene pairs, stand-alone sensor domain genes, and c-di-GMP turnover genes (Fig. 2b and Extended Fig. 3). For example, the complexity apex of TCSs in *Campylobacterota* is represented by *Sulfurimonas gotlandica* GD1, with the total number of solo HKs, HHKs and HRRs in combination reaching 45% (Fig. 2b and Supplementary Table 3). This genome has 10 clusters enriched in atypical HKs; the biggest cluster contains 22 signaling proteins (Fig. 2d and Extended Fig. 4). Functional relatedness suggests that proteins encoded by the same gene cluster might participate in the same pathway(s) with several sensors and checkpoints, organized as a branched and multi-step phosphorelay to elicit a specific output response. Moreover, REC domain is the most recurrent module in these clusters, either fused with other modules to link different transmitters or standing alone as the phosphate sink to fine-tune the signaling process (Extended Fig. 4).

Although the abundance of TCSs can be quantified, their evolutionary modes cannot be precisely measured. We found that fusion of duplication and HGT pieces produces many chimeric HKs and RRs with non-DNA-binding domain outputs. Meanwhile, it is challenging to distinguish between vertical inheritance after an “earlier” HGT event with many gene losses and the recent HGTs among closely related species, especially with the avalanche of genomic data nowadays. In spite of these difficulties in quantification of gene duplication and HGT ratios, it is generally accepted that both of them provide the raw materials to expand TCS pathways. During bacterial niche expansion, massive gene duplications and HGTs might easily result in new domain organizations of signaling proteins that are highly modular. Thus, more atypical HKs including HHKs and HRRs, and also more atypical RRs with non-DNA-binding domain outputs emerge while the TCS size increases, leading to the *Matthew Effect* in signaling pathways. Besides, our results suggest that REC domains promote connectivity as well as genetic innovation to increase the complexity of TCS.

### C-di-GMP is made for connection

C-di-GMP is the most ubiquitous secondary messenger in bacteria. It is produced from two GTP molecules by diguanylate cyclases (DGCs) with GGDEF domain, and is broken down by phospodiesterases (PDEs) that contain either EAL or HD-GYP domain ^40^. The core of c-di-GMP signaling system is composed of enzymes that make and break it and diverse effectors that it can bind to regulate biofilm, motility, cell cycle, virulence, etc ^41^. The complexity of this system has long been realized but not yet understood, owing to the inherent “many to many” links connected by this single molecule (Fig. 3a) ^42^. Global c-di-GMP regulation mode consists of input signal integration and output divergence, based on the diffusible nature of this molecule and the fact of “more enzymes, less effectors” identified in many species. In contrast, local c-di-GMP signaling pathways employ physical interaction or complex formation of specific DGC, PDE, effector and target to ensure specificity. Moreover, both local and global regulation can form dynamic links with spatiotemporal control ^43,44^.

**Fig. 3.**
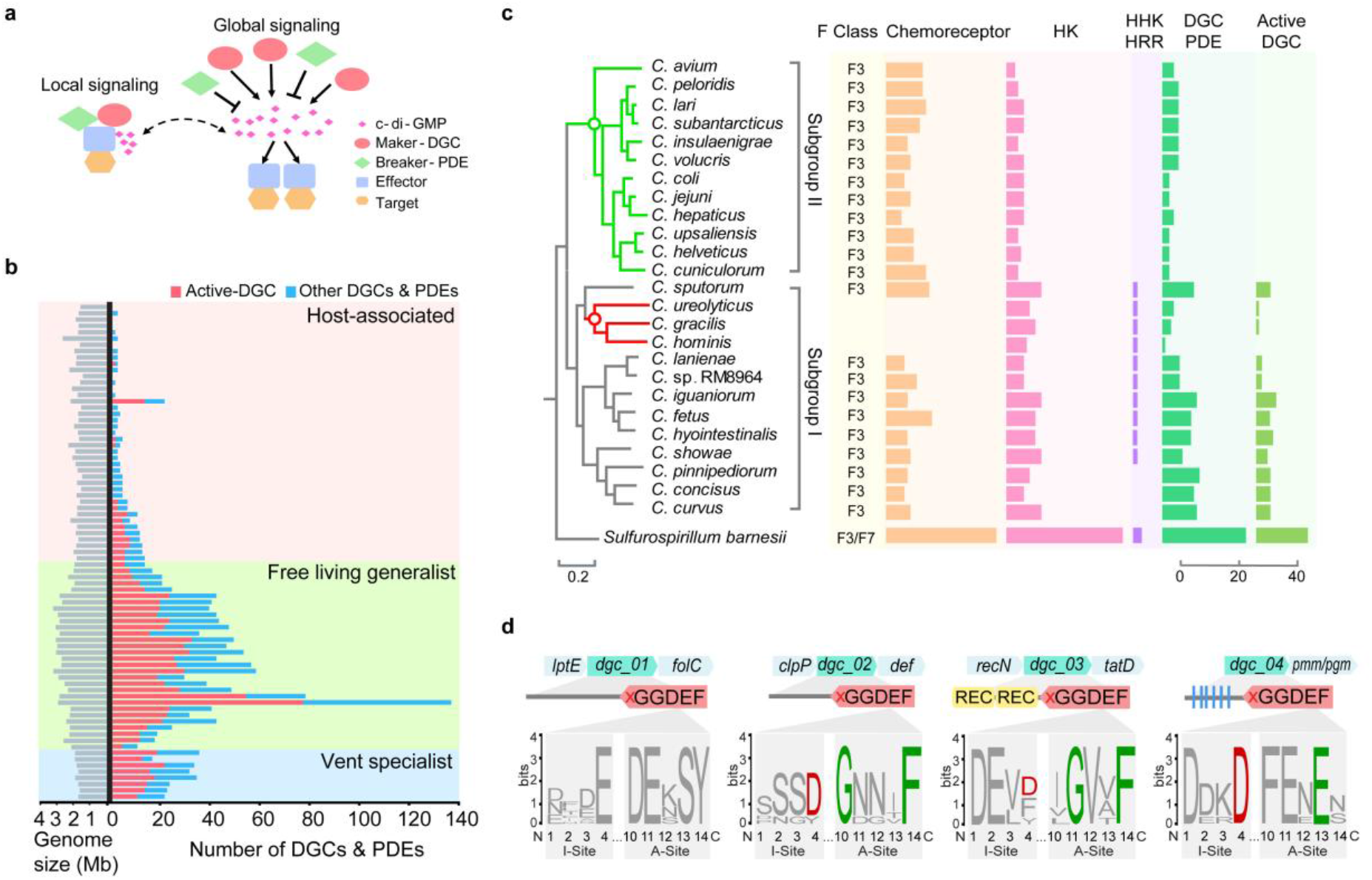
c-di-GMP mediated system in *Campylobacterota*. **(a)** Local and global c-di-GMP signaling system. **(*b*)** The abundance of DGCs and PDEs in *Campylobacterota* species. The category and number of DGCs and PDEs in each genome in Supplementary Table 4. **(c)** The number of signaling proteins in the *Campylobacter* genus. The phylogenetic tree is taken from Extended Fig. 1. In the tree, green circle represents the branch point of subgroup II when the c-di-GMP signaling system is lost; red circle represents the branch point of a lineage losing flagellar genes. **(*d*)** Four inactive DGCs in species of subgroup II in Fig. 3c. The flanking genes of these 4 DGCs are indicated above and their domain organization, degenerate enzymatic (A-site) and c-di-GMP binding (I-site) motifs are shown below. Active I-site is generally featured by RxxD, and the active A-site is characterized by GGD/EEF, so the GGDEF domain of the 4 DGCs here are marked by red cross implying enzymatically inactive.

Due to the lack of knowledge of all the effectors, we assessed the complexity level of c-di-GMP mediated system by the quantification of its turnover enzymes. DGCs and PDEs can be readily identified by their dedicated domains (GGDEF, EAL or HD-GYP domains). Our domain analysis of all DGCs and PDEs revealed that 98% are multi-module, with N-terminal sensory domains in common with chemoreceptors and HKs, or with REC domains mostly having cognate HKs in close proximity (Extended Fig. 5). Thus, the number of c-di-GMP enzymes can represent the sensing ability of the species. In addition, for species with multiple c-di-GMP enzymes, their sensory domains or connected sensory proteins (such as HKs) are different, thus no functional redundancy exists for these enzymes (Extended Fig. 5). However, not all identified DGCs and PDEs are active, and enzymatically inactive GGDEF or EAL domains often serve as c-di-GMP binding effectors ^45^. If no active DGC is encoded in the genome, no c-di-GMP can be produced to mediate signal transduction, even in the presence of degenerate domains. Hereafter, we separate the proteins with enzymatically active GGDEF domain called “active DGCs” from all the other proteins containing EAL, HD-GYP or degenerate GGDEF domains as the rest of DGCs and PDEs (Supplementary Table 4).

The total number of DGCs and PDEs within *Campylobacterota* species varies greatly, ranging from 0 to 137, indicating that bacterial genomes gain and lose c-di-GMP enzymes with ease (Fig. 3b). Vent specialists generally have >20 c-di-GMP turnover enzymes, more than their average number of HKs or chemoreceptors, implying that c-di-GMP signaling pathways play an important role in ancient lineages. Free-living generalists gain more enzymes, with upsurge in two species but no big increase of their genome size (Fig. 3b and Supplementary Table 4). This extreme proliferation of DGCs and PDEs is accompanied by F3/F7/F9 chemosensory classes and a larger number of TCS genes in these two species (Fig. 1a and 2b), indicative of many gene gain events of signaling systems *en masse* during niche adaptation. The sheer abundance of DGCs and PDEs drops abruptly in host-associated species (Fig. 3b). No active DGC could be found in the *Helicobacter* genus except for one species and more than half of species of the *Campylobacter* genus, meaning that c-di-GMP signaling system had disappeared in these host-associated species (Supplementary Table 4). Overall, the number of c-di-GMP enzymes in most *Campylobacterota* species correlates with their ecotype range (Fig. 3b).

The ubiquity of c-di-GMP enzymes within *Campylobacterota* suggests that c-di-GMP signaling system evolved in the ancestor of this phylum and greatly expanded in free-living generalists during niche expansion. The upsurge of c-di-GMP enzymes to 137 in *S. gotlandica* GD1 compared to 30∼40 in other species of the same genus raises an interesting question: How does a bacterium quickly gain so many gene copies of c-di-GMP enzymes? Sequence analyses of all 137 c-di-GMP enzymes revealed only two pairs of proteins with identical domain organization and high sequence similarity, suggestive of recent gene duplications (Extended Fig. 5). Other enzymes with highly similar GGDEF domains have different types of sensory domains or show diverse combination of sensory domains, GGDEF and EAL domains, unlikely produced by single gene duplication event (Extended Fig. 5). Thus, the upsurge of c-di-GMP enzymes in bacterial species might be achieved by mixed evolutionary modes: both HGTs and gene duplications provide the genetic material, and gene fusion/fission and fast sequence divergence contribute to make new genes.

Another intriguing question is the absence of c-di-GMP signaling system in host-associated pathogens such as *Helicobacter pylori* and *Campylobacter jejuni*, since c-di-GMP plays an important role in regulation of virulence in many other pathogens. To approach the determinants of c-di-GMP signaling pathways, we examined the gradual change of all signaling proteins for the *Campylobacter* genus based on our “eco-evo” framework. All *Campylobacter* species have greatly reduced numbers of components for all signal transduction systems compared to its close relative *Sulfurospirillum* genus, suggesting that substantial gene loss of signaling proteins happened for the last common ancestor of the *Campylobacter* genus, likely due to the association with animal hosts (Fig. 3c). Within this genus, two distinctive clusters can be defined: Subgroup I with relatively more signaling proteins such as TCSs and c-di-GMP enzymes and Subgroup II with less signaling proteins and no canonical active DGCs. To verify the absence of active DGCs in Subgroup II, we further examined 443 strains of *Campylobacter* species with completely sequenced genomes (Supplementary Table 5). For Subgroup II, all the genomes still encode 1∼4 proteins with degenerate GGDEF domains and do not possess an active GGDEF domain, or EAL/HD-GYP domains (Supplementary Table 5). Importantly, these degenerate GGDEF domains within Subgroup II also have degenerate inhibitory sites for c-di-GMP binding, indicating that they are unlikely to serve as c-di-GMP effectors, either (Fig. 3d). Furthermore, the species of subgroup II only retain 4∼6 TCS pairs but no HHK or HRR, one F3 chemosensory class, only one STYK found in one species, no cAMP medicated pathways or ECFs (Fig. 3c). Except for the single chemosensory class for flagellar motility, other signal transduction systems in these species are basically OCS proteins and very few TCS pairs, lacking components and links compared to the subgroup I with active DGCs. Hence, it is possible that the presence of c-di-GMP signaling system requires a minimal signal transduction network size or connectivity to build more links into it.

### The evolution of other signaling systems in *Campylobacterota*

ECFs are typically regulated by their cognate anti-σ factors that can sense environmental stimuli, and they guide the RNA polymerase to initiate expression of genes related to stress responses ^46^. Recent census studies of ECF proteins greatly expanded their known diversity and re-classified them into 157 groups ^18^. However, only 2 ECF groups were identified in several *Campylobacterota* species. One ECF group, ECF238 (also called SigZ), is found in just one *Arcobacter* species acquired through HGT from δ-proteobacteria; the other group, ECF242, is present in 3 genera including *Arcobacter, Sulfurospirillum*, and *Wolinella* (Fig. 4a and Supplementary Table 6). The presence of ECFs in these later evolved genera suggests that the ECF system was introduced into *Campylobacterota* during niche expansion by HGT. ECF242 is composed of a σ factor FecI, an anti-σ factor FecR and a TonB-dependent receptor FecA ^18^. Only half of the species of the above 3 genera have ECF242 gene cluster and most of them contain 2 or more copies (Fig. 4a). Phylogenetic analyses of FecI and FecR protein do not show the same topology as the species tree, indicating that HGT events happened multiple times among these 3 genera alongside gene duplications (Extended Fig. 6). Nevertheless, FecI and FecR protein trees share the same clustering pattern, supporting the co-evolution of an ECF σ factor and its anti-σ factor.

**Fig. 4.**
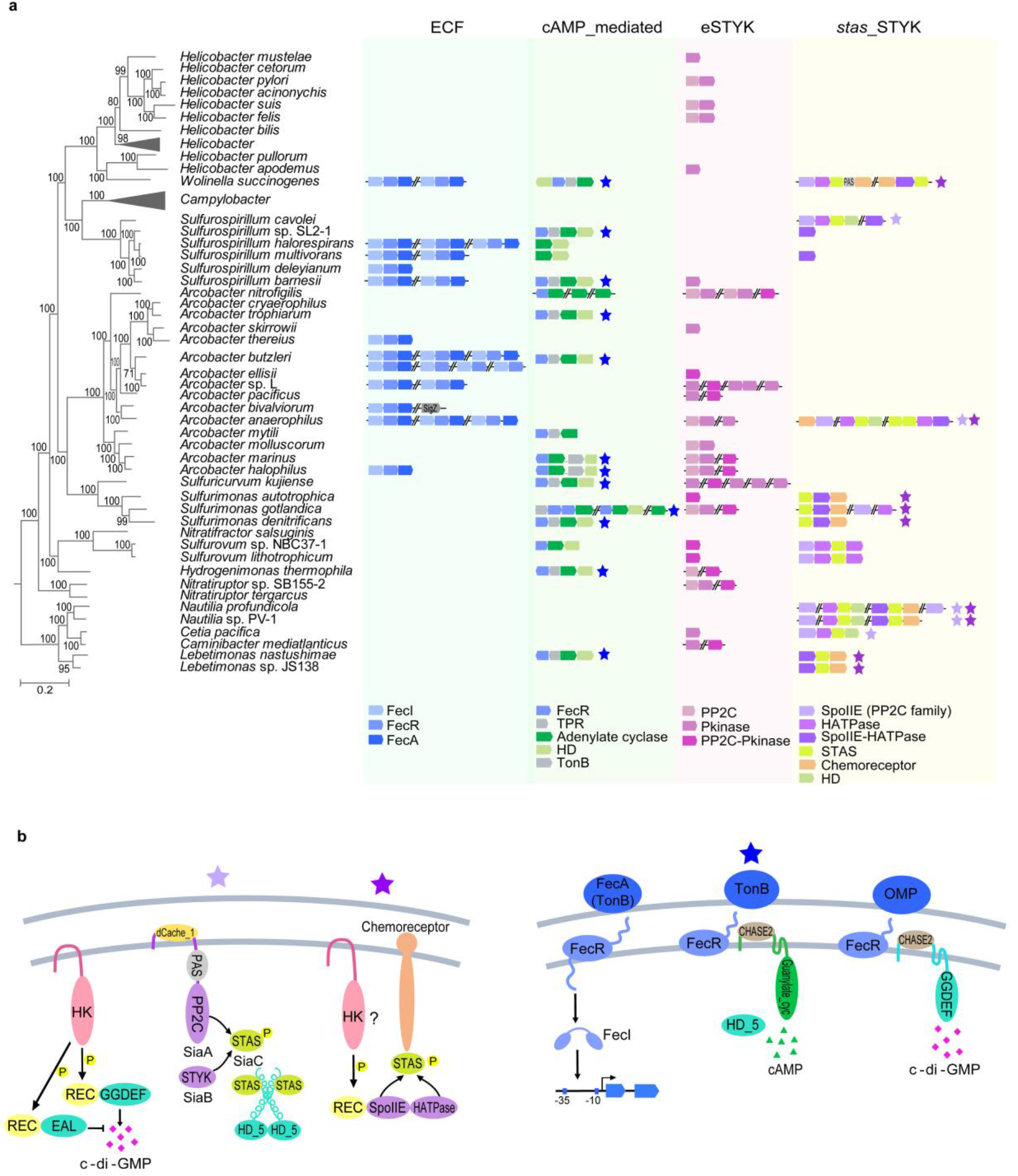
The ECF, cAMP, STYK systems in *Campylobacterota*. **(*a*)** Phylogenetic profile of ECF, cAMP, STYK systems in *Campylobacterota*. Genes or gene clusters constituting the ECF, cAMP or STYK systems are shown for species that contain them. The *Campylobacter* genus and partial *Helicobacter* species that do not have ECF, cAMP or STYK signaling systems are collapsed. SpoIIE protein or SpoIIE domain that dephosphorylate STAS protein belong to PP2C family. **(*b*)** Potential pathways with combination of two or more systems. Left: Examples of potential pathways found in *Nautilia profundicola* AmH; Right: classical ECF pathway and ECF components potentially interacting with cAMP or c-di-GMP enzymes in *Arcobacter halophilus* CCUG 53805. Stars with different color in b are also labeled in a, meaning the presence of the potential pathways in the species.

cAMP is produced by adenylate cyclase and binds cAMP receptor protein (CRP) or few other proteins to control diverse cellular functions ^47^. Only 21% of *Campylobacterota* species encode adenylate cyclase, but 86% species have one or more CRP-like proteins, indicating that CRP homologs in the species without adenylate cyclase are not likely to be regulated by cAMP (Supplementary Table 6). Notably, all adenylate cyclases identified in *Campylobacterota* species have the same sensory domain (CHASE2 domain), and most of them are located in a cluster containing a phosphodiesterase that can hydrolyze cAMP, a FecR domain-containing outer membrane protein and a TPR domain-containing protein (Fig. 4a). The clustering of these four genes in multiple *Campylobacterota* species suggests that they likely participate in the same signal transduction pathway. Phylogenetic analysis of the conserved adenylate cyclase did not yield same branch pattern as the species tree (Extended Fig. 7), indicative of HGT events of this gene cluster among free-living *Campylobacterota* species as well as other bacterial phyla.

STYKs that can phosphorylate serine, threonine, or tyrosine residues in target proteins have two kinds in bacteria and they are evolutionarily unrelated ^48^. One kind are eukaryotic-type protein kinases (eSTYKs) characterized by the presence of Pfam’s Pkinase domain (PF00069); the other kind are members of Histidine kinase-like ATPase family but they phosphorylate serine/threonine residues in specific proteins with a single STAS domain (also called anti-sigma-factor antagonists). Here, we name the 2^nd^ kind of STYKs with HATPase domain but no HisKA domain (thus not HK of TCS) as *stas*_STYKs. Both eSTYKs and *stas*_STYKs share the same partner phosphatase, PP2C family phosphatase, to reverse the phosphorylation reaction. One third of *Campylobacterota* species have eSTYKs, either an eSTYK and a cognate PP2C phosphatase, or a bifunctional kinase/phosphatase fusing PP2C and Pkinase domains together (Fig. 4a and Supplementary Table 6). Only one fifth of the *Campylobacterota* species have *stas*_STYKs genes, mostly having target *stas* and cognate PP2C phosphatase genes in the same neighborhood in the genome (Fig. 4 and Supplementary Table 6). Except for their clustering pattern with cognate phosphatases, the species distribution of both eSTYKs and *stas*_STYKs in *Campylobacterota* is sporadic, apparently resulting from multiple HGT events. For example, the eSTYKs and their phosphatase genes in 7 *Arcobacter* species reside in the middle of Type VI secretion system loci, which are prone to HGT (Supplementary Table 6).

Taken together, in contrast to the prevalence of chemosensory, TCS and c-di-GMP systems in *Campylobacterota* species, ECF, cAMP and STYK systems are mostly present in free-living generalists with sporadic distribution. Our phylogenetic analyses suggest that the last common ancestor of *Campylobacterota* already evolved F3 chemosensory class, TCS and c-di-GMP systems, thus vertical inheritance, gene duplication and HGT can greatly expand these ancestral systems and result in increased complexity. However, ECF, cAMP and STYK systems apparently were acquired multiple times by HGT, and were mostly lost in host-associated *Campylobacter* and *Helicobacter*. Aside from the 6 signaling systems described above, the *Campylobacterota* phylum does not have c-di-AMP and QS systems. It should be mentioned that QS differs from all other signal transduction systems in terms of sensing self-derived rather than environmental signals. Although none of the known QS pathways has been identified in any *Campylobacterota* species, our domain analyses of known QS components in model organisms from other phyla indicate that QS receptors belong to OCS, TCS, c-di-GMP enzymes or chemoreceptors (Extended Fig. 8). Thus, we conclude that QS adopts existing modules to merge into the signaling network without *de novo* innovation in transmitter type or mechanism.

### Complexity beyond each signaling system

Theoretically, the modularity of signaling proteins enables the merge of different mechanisms into one pathway, and bacteria had already used this strategy to assemble multi-step information processing during evolution. As described earlier, TCSs are frequently merged with other systems through the fusion of REC domain with non-canonical output domains. In addition, one of the chemosensory classes in *Pseudomonas aeruginosa* is reported to regulate c-di-GMP concentration because its response regulator is fused with GGDEF domain ^49^. Our analyses also discovered potential pathways combined with two or more systems, based on neighborhood gene cluster, known protein-protein or domain-domain interactions, and signaling flow logic from sensors to transmitters (Fig. 4b). For example, based on their co-occurrence in multiple *Campylobacterota* species, a PP2C phosphatase with periplasmic sensory domain and a *stas*_STYK probably regulate a c-di-GMP turnover enzyme by the phosphorylation and dephosphorylation of a STAS protein. Similarly, a bifunctional PP2C-STYK protein with N-terminal REC domain that is likely regulated by an unknown HK protein, and itself might affect a chemoreceptor conformation by its target STAS protein (Fig. 4a,b and Supplementary Table 6). These potential pathways in *Campylobacterota* species lack experimental verification so far, but the merge of STYK-PP2C and c-di-GMP systems has been recently reported as SiaABCD pathway in *P. aeruginosa* ^50,51^. This pathway is an example of a c-di-GMP enzyme (SiaD) regulated by PP2C (SiaA), *stas*_STYK (SiaB) and STAS protein (SiaC). Another recent study proposed a working model for ECF modules for inhibition of transmembrane cAMP enzyme activity ^52^. Our analyses revealed clustering of genes encoding ECF sensors, partial ECF transmitters and transmembrane cAMP or c-di-GMP enzymes in multiple *Campylobacterota* species, suggesting their functional relationship (Fig. 4a,b and Supplementary Table 6).The common theme of these potential pathways is to convert extracellular signals to intracellular secondary messengers or protein phosphorylation-based transmission processes, perhaps allowing for multiple input signal integration and output divergence (Fig. 4b). Future experimental validations are needed for these pathways with combined systems, which will provide insights for the development of sophisticated genetic circuit in synthetic biology.

## DISCUSSION

Here, we defined the complexity of each system differently based on their signal transduction mechanism, which determines how basic interactions among components are formed and more interactions can be propagated. Thus, we cannot simply group signal transduction systems such as OCS, TCS and others, but rather recognize their distinctive mechanisms and then disentangle the network evolution. In the “eco-evo” context of the *Campylobacterota* phylum, we compared the dynamic changes of all signaling systems and revealed that bacterial signaling capacity and complexity arise and drop together.

Niche expansion of the species leads to a larger genetic pool from the microbial community which facilitates HGT; meanwhile, stresses encountered along environmental changes may cause mistakes during DNA replication followed by repair, resulting in gene duplication, loss, fusion and fission. These genetic events not only propagate existing protein interactions, but also create opportunities for innovation of new genes/pathways, and combination of different mechanisms in one signaling pathway. Thus, free-living generalists with large signaling networks tend to employ multi-step or branched pathways to achieve tunable information processes (Fig. 5). On the contrary, when species become specialized in a restricted habitat, a large sensory repertoire is no longer required but can be energy costive. Thus, signaling networks shrink together with their regulated targets or pathways, leading to a streamlined genome for specialist lifestyle. Moreover, host-associated specialists generally retain simple OCSs and classical TCSs, but lose complex pathways with multi-components and many links (Fig. 5). The gradual disappearance of c-di-GMP system and other systems in the *Campylobacter* genus provide such an example. However, if the signal outputs are retained in specialists, their co-evolved signaling systems are still needed. For example, both *H. pylori* and *C. jejuni*, with very streamlined genomes and minimal signaling network size, still retain multi-component chemosensory system to control their flagellar motility (Fig. 5). Thus, host-associated specialists tend to keep their signal transduction simple and straightforward in a small network, but they often show biased preference for signaling systems determined by co-evolution with the outputs.

**Fig. 5.**
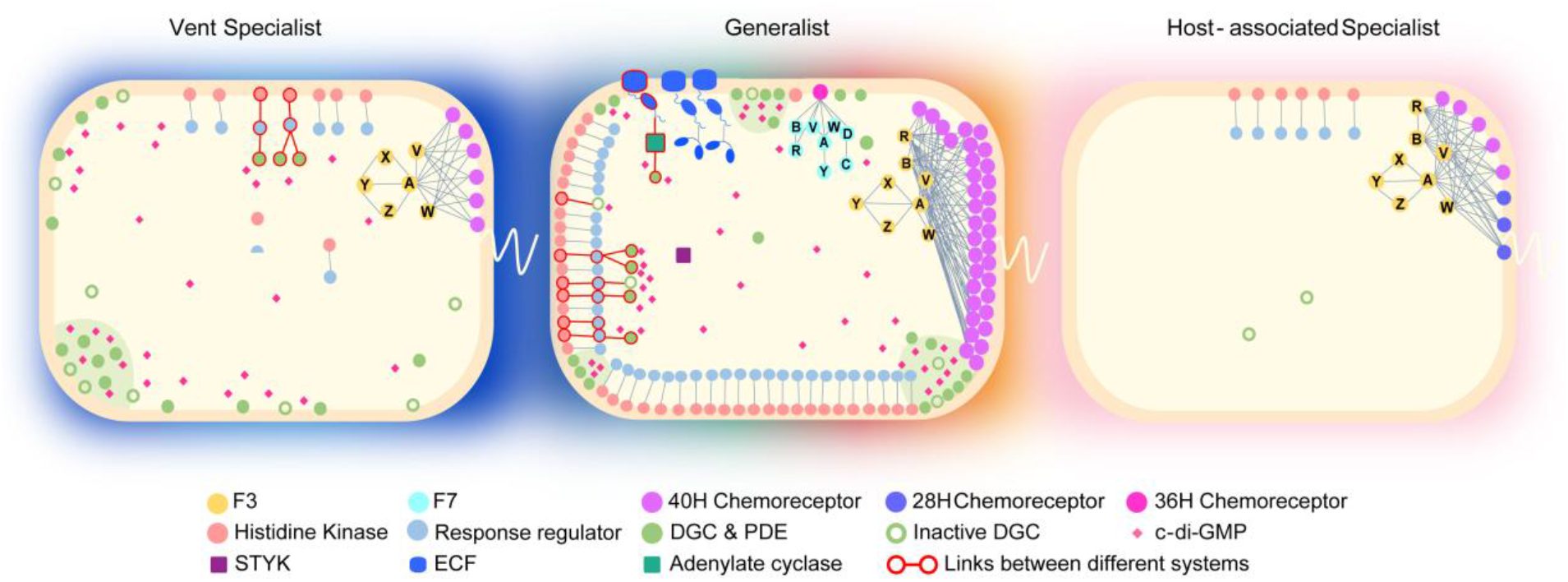
Signaling networks in representative species from diverse ecological niches. Vent specialist: *Lebetimonas natsushimae* HS1857; Generalist: *Sulfurospirillum barnesii* SES-3; Host-associated specialist: *Campylobacter jejuni* 81-176. The shading color around the cell depicts the ecotype range of the representative species: Blue, deep-sea hydrothermal vent; Iridescence, various environments; Pink, host-associated. Transmembrane DGCs and PDEs are arbitrarily clustered within cell to represent local c-di-GMP signaling.

All links in the signaling network are based on protein-protein or protein-ligand interactions. To ensure signaling specificity, co-evolution of interactive partners is required to avoid nonspecific crosstalk, which is best exemplified by classical TCSs ^53,54^. Actually, co-evolution exists at all protein-protein interactions in all of the signal transduction systems, including chemosensory classes, c-di-GMP local signaling complexes, ECF σ factors and anti-σ factors, *stas*_STYKs and STASs. Co-evolution constraints normally apply to the interaction interfaces of components within one pathway. If many interactions are involved in signal transmission, many proteins need to co-evolve, and as a result, their evolutionary flexibility will decrease and vice versa. Hence, multi-component chemosensory system generally adds new pathways by acquiring entire chemosensory classes through HGT, but simpler signaling systems with two or few components can gain more transmitters by both gene duplication/divergence and HGT. However, secondary messenger such as c-di-GMP can break the co-evolution constraint if the link between an enzyme and an effector solely depends on this same molecule with no direct protein-protein interactions involved. Due to this flexibility, a c-di-GMP enzyme gained by gene duplication or HGT can directly generate links with existing effectors. Likewise, if a c-di-GMP enzyme loses its function, its effector(s) can be easily rewired by other existing enzymes to complement its function under selection, as demonstrated by laboratory evolution experiments ^55,56^. This might explains why bacterial species with c-di-GMP mediated system gain and lose c-di-GMP enzymes very dynamically, compared to other signaling systems.

Finally, it is extremely difficult to delineate the signaling pathways for a bacterial species with dozens of signaling homologs at its disposal. Our protein domain analyses suggest that the activities of transmitter domains are directly or indirectly regulated by different sensory domains responding to diverse stimuli. Hence, no functional redundancies really exist for signaling homologs in one bacterial cell with a complex network, but puzzle for us to decipher and then apply in genetic circuit design.

## METHODS

### Data sources

The statistics of completely sequenced genomes of various bacterial phyla were collected in NCBI RefSeq database and the taxonomic ranks for some new phyla were updated according to GTDB database (Supplementary Table 1) ^24,25^. For *Campylobacterota* species with multiple completely sequenced strains, only one strain was selected to represent the species. To cover the most species diversity and also ensure genome quality, 74 complete genomes and 8 incomplete but close to complete genomes were included for analyses (Supplementary Table 2). All these 82 genome sequences were downloaded from NCBI genome database (https://www.ncbi.nlm.nih.gov/genome/microbes/). The ecological and physiological information of all strains were extracted from references listed in Supplementary Table 2. For analyses of c-di-GMP enzymes in the *Campylobacter* and *Helicobacter* genera, all completely sequenced genomes from both genera were downloaded from the same NCBI genome database. The numbers of complete genomes for each *Campylobacter* or *Helicobacter* species were summarized in Supplementary Table 5.

### Bioinformatics software

Phylogenetic trees are constructed either by FastTree 2.0 or Mega X programs as indicated below ^57,58^. All homologous protein searches were conducted by blast-2.12.0 ^59^; multiple sequence alignments were performed by MAFFT program using the e-ins-i algorithm ^60^; domain architectures of all proteins were identified by SMART and HHpred ^61,62^. TreeCollapse was used to produce the final tree figure in Fig. 1d with large dataset ^63^. Sequence logos for multiple sequence alignments in Fig. 3d were generated by Weblogo ^64^

### Phylogenomic tree construction for *Campylobacterota*

82 genomes from *Campylobacterota* and 6 genomes from closely related *Desulfurellales* class as the out-group were retrieved to reconstruct the species tree for the *Campylobacterota* phylum. Phylogenetic inference was performed using the UBCG pipeline ^65^. Basically, 92 single-copy orthologous genes were identified from these genomes by Prodigal and HMMER with default values ^66,67^. Orthologous protein sequences were aligned by MAFFT and the alignment positions with gap characters from more than 50% sequences were excluded by Gblocks ^68^. A maximum likelihood tree based on concatenated protein sequence alignments was constructed in FastTree using JTT + CAT model, and bootstrap analysis was carried out using 1000 replications.

### Analyses of chemosensory system

All protein components of the chemosensory system (CheA, CheW, CheV, CheB, CheR, CheC, CheD, CheX, CheY, CheZ and Chemoreceptors) were identified by blast-2.12.0 against 82 *Campylobacterota* genomes. The blast hits with e-value less than e-5 were assigned as potential homologs and all the candidates were re-examined by searching at MiST, SMART and HHpred databases manually ^19^. The chemosensory genes were mapped on the linearized genomes and visualized by RIdeogram ^69^. The chemosensory F class were assigned based on ref ^30^. The H classes for all chemoreceptors were manually assigned based on ref ^70^, and also verified using MiST database ^19^.

A maximum-likelihood phylogenetic tree was constructed based on concatenated sequence alignments of CheV, CheA and CheW of F3 class from *Campylobacterota* genomes. The tree was constructed in MEGA X using WAG model with bootstrap 100 replications. Another phylogenetic tree based on 2,136 CheA homologs of different F classes from various bacterial phyla (1,354 species) was constructed by FastTree using JTT + CAT model.

### Analyses of TCS

Both HisKA (Pfam’s no: PF00512) and HATPase (PF02518) domains are used as query sequence for searching HKs, and REC domain (PF00072) as the query for RRs in each genome by blast-2.12.0. In addition, if a REC domain is found at the C-terminus after or N-terminus before both HisKA and HATPase domains in a protein sequence, it was considered as HHK or HRR, respectively. A solo HK or HHK gene is more than four genes away from a RR gene. The classification of TCS proteins in each genome was presented in Supplementary Table 3. The gene neighborhood of TCS genes was manually checked in selected species with abundant HHK and HRR genes. All signal transduction genes were plotted on the linearized genomes of these species by RIdeogram. The domain architectures of all HKs and RRs in representative species of each genus was shown in Extended Data Fig. 4.

### Analyses of DGCs and PDEs

We identified all DGCs and PDEs by blast searching GGDEF (PF00990), EAL (PF00563) and HD-GYP (PF13487) domains against all 82 *Campylobacterota* genomes, according to the characteristics of DGCs and PDEs summarized by ref ^40^. Protein domain architecture was analyzed by SMART. DGCs with active GGDEF domain should have GGDEF/GGEEF/SGDEF/AGDEF motif at the A-site, otherwise were considered as degenerate DGCs. We classified all the DGCs and PDEs into 6 categories in each species: total DGCs; DGCs with partner domain (potential sensory domain or domain mediating interaction with another protein); PDEs with partner domain; bifunctional GGDEF-EAL/HD-GYP domain-containing proteins; single domain DGCs and PDEs; DGCs with active GGDEF domain (Supplementary Table 4). The domain architectures of DGCs and PDEs in representative species of each genus was shown in Extended Data Fig. 5.

### Analyses of ECF, cAMP and STYK systems

ECF σ factors were identified using Sigma 70_r2 (PF04542) and Sigma70_r4_2 (PF08281) domains as query for blast search, and only blast hits with both and only these two domains were counted as ECF σ factors according to ref ^18^. The gene neighborhood was manually analyzed to determine the ECF groups according to ref ^18^. Phylogenetic tree of FecI and FecR proteins in *Campylobacterota* were constructed by Mega X following the methods above.

Both Adenylate_cycl (PF01295) and Guanylate_cyc (PF00211) domains was used as query for blast search of adenylate cyclase in each genome. Phylogenetic tree of adenylate cyclases in *Campylobacterota* were constructed following the method above and the gene neigbhorhood was manually checked.

For identification of eSTYKs systems, the Pkinase (PF00069) and PP2C (PF00481) domains was used as query to perform blast search. For *stas*_STYKs systems, the SpoIIE (PF07228) and HATPase (PF13581) domains were used as query.

All the protein ID of the above 3 systems were summarized in Supplementary Table 6.

### Analyses of QS pathways

The enzymes and receptors of six well studied QS pathways in representative strains were summarized from previous studies in Extended Data Fig. 8. The domain structure of both synthetases and receptors were identified using SMART. The protein sequences of synthetases and the potential ligand binding domains of the receptors were used as probe to do blast search against *Campylobacterota* genomes. None of the genomes return both hits for synthetase and corresponding receptor of each QS pathway. Thus, we conclude that none of the QS pathways is present in analyzed genomes of the *Campylobacterota* phylum.

## Acknowledgements

The authors would like to thank Drs. Michael Galperin and Wei Qian for critical reading and comments of the manuscript. This research was supported by National Natural Science Foundation of China (31870064), Key Special Project for Introduced Talents Team of Southern Marine Science and Engineering Guangdong Laboratory (Guangzhou) (GML2019ZD0407), Strategic Priority Research Program of the Chinese Academy of Sciences (XDA19060301), and Innovation Academy of South China Sea Ecology and Environmental Engineering, Chinese Academy of Sciences (NO.ISEE2021ZD03).

## Author contributions

BG designed research; RM, YL, YC and YM performed research; BG and RM analyzed data and wrote the paper.

## REFERENCES

1 Adler, J. Chemotaxis in Escherichia coli. Cold Spring Harb Symp Quant Biol 30, 289–292, doi:10.1101/sqb.1965.030.01.030 (1965).

2 Ninfa, A. J. & Magasanik, B. Covalent modification of the glnG product, NRI, by the glnL product, NRII, regulates the transcription of the glnALG operon in Escherichia coli. Proceedings of the National Academy of Sciences of the United States of America 83, 5909–5913, doi:10.1073/pnas.83.16.5909 (1986).

3 Nixon, B. T., Ronson, C. W. & Ausubel, F. M. Two-component regulatory systems responsive to environmental stimuli share strongly conserved domains with the nitrogen assimilation regulatory genes ntrB and ntrC. Proceedings of the National Academy of Sciences of the United States of America 83, 7850–7854, doi:10.1073/pnas.83.20.7850 (1986).

4 Makman, R. S. & Sutherland, E. W. Adenosine 3’,5’-Phosphate in Escherichia Coli. The Journal of biological chemistry 240, 1309–1314 (1965).

5 Ross, P. et al. Regulation of cellulose synthesis in Acetobacter xylinum by cyclic diguanylic acid. Nature 325, 279–281, doi:10.1038/325279a0 (1987).

6 Witte, G., Hartung, S., Buttner, K. & Hopfner, K. P. Structural biochemistry of a bacterial checkpoint protein reveals diadenylate cyclase activity regulated by DNA recombination intermediates. Mol Cell 30, 167–178, doi:10.1016/j.molcel.2008.02.020 (2008).

7 Davies, B. W., Bogard, R. W., Young, T. S. & Mekalanos, J. J. Coordinated regulation of accessory genetic elements produces cyclic di-nucleotides for V. cholerae virulence. Cell 149, 358–370, doi:10.1016/j.cell.2012.01.053 (2012).

8 Whiteley, A. T. et al. Bacterial cGAS-like enzymes synthesize diverse nucleotide signals. Nature 567, 194–199, doi:10.1038/s41586-019-0953-5 (2019).

9 Munoz-Dorado, J., Inouye, S. & Inouye, M. A gene encoding a protein serine/threonine kinase is required for normal development of M. xanthus, a gram-negative bacterium. Cell 67, 995–1006, doi:10.1016/0092-8674(91)90372-6 (1991).

10 Lonetto, M. A., Brown, K. L., Rudd, K. E. & Buttner, M. J. Analysis of the Streptomyces coelicolor sigE gene reveals the existence of a subfamily of eubacterial RNA polymerase sigma factors involved in the regulation of extracytoplasmic functions. Proceedings of the National Academy of Sciences of the United States of America 91, 7573–7577, doi:10.1073/pnas.91.16.7573 (1994).

11 Fuqua, W. C., Winans, S. C. & Greenberg, E. P. Quorum Sensing in Bacteria - the Luxr-Luxi Family of Cell Density-Responsive Transcriptional Regulators. Journal of bacteriology 176, 269–275 (1994).

12 Ulrich, L. E., Koonin, E. V. & Zhulin, I. B. One-component systems dominate signal transduction in prokaryotes. Trends in microbiology 13, 52–56, doi:10.1016/j.tim.2004.12.006 (2005).

13 Galperin, M. Y. Bacterial signal transduction network in a genomic perspective. Environmental microbiology 6, 552–567, doi:10.1111/j.1462-2920.2004.00633.x (2004).

14 Galperin, M. Y. A census of membrane-bound and intracellular signal transduction proteins in bacteria: bacterial IQ, extroverts and introverts. BMC microbiology 5, 35, doi:10.1186/1471-2180-5-35 (2005).

15 Galperin, M. Y. What bacteria want. Environmental microbiology 20, 4221–4229, doi:10.1111/1462-2920.14398 (2018).

16 Alm, E., Huang, K. & Arkin, A. The evolution of two-component systems in bacteria reveals different strategies for niche adaptation. PLoS Comput Biol 2, e143, doi:10.1371/journal.pcbi.0020143 (2006).

17 Seshasayee, A. S., Fraser, G. M. & Luscombe, N. M. Comparative genomics of cyclic-di-GMP signalling in bacteria: post-translational regulation and catalytic activity. Nucleic acids research 38, 5970–5981, doi:10.1093/nar/gkq382 (2010).

18 Casas-Pastor, D. et al. Expansion and re-classification of the extracytoplasmic function (ECF) sigma factor family. Nucleic acids research 49, 986–1005, doi:10.1093/nar/gkaa1229 (2021).

19 Gumerov, V. M., Ortega, D. R., Adebali, O., Ulrich, L. E. & Zhulin, I. B. MiST 3.0: an updated microbial signal transduction database with an emphasis on chemosensory systems. Nucleic acids research 48, D459–D464, doi:10.1093/nar/gkz988 (2020).

20 Galperin, M. Y., Higdon, R. & Kolker, E. Interplay of heritage and habitat in the distribution of bacterial signal transduction systems. Mol Biosyst 6, 721–728, doi:10.1039/b908047c (2010).

21 Gouy, R., Baurain, D. & Philippe, H. Rooting the tree of life: the phylogenetic jury is still out. Philos Trans R Soc Lond B Biol Sci 370, 20140329, doi:10.1098/rstb.2014.0329 (2015).

22 Coleman, G. A. et al. A rooted phylogeny resolves early bacterial evolution. Science 372, doi:10.1126/science.abe0511 (2021).

23 Waite, D. W. et al. Comparative Genomic Analysis of the Class Epsilonproteobacteria and Proposed Reclassification to Epsilonbacteraeota (phyl. nov.). Front Microbiol 8, 682, doi:10.3389/fmicb.2017.00682 (2017).

24 Schoch, C. L. et al. NCBI Taxonomy: a comprehensive update on curation, resources and tools. Database (Oxford) 2020, doi:10.1093/database/baaa062 (2020).

25 Parks, D. H. et al. A standardized bacterial taxonomy based on genome phylogeny substantially revises the tree of life. Nature biotechnology 36, 996–1004, doi:10.1038/nbt.4229 (2018).

26 Campbell, B. J., Engel, A. S., Porter, M. L. & Takai, K. The versatile epsilon-proteobacteria: key players in sulphidic habitats. Nature reviews. Microbiology 4, 458–468, doi:10.1038/nrmicro1414 (2006).

27 Li, M. & Hazelbauer, G. L. Core unit of chemotaxis signaling complexes. Proceedings of the National Academy of Sciences of the United States of America 108, 9390–9395, doi:10.1073/pnas.1104824108 (2011).

28 Gumerov, V. M., Andrianova, E. P. & Zhulin, I. B. Diversity of bacterial chemosensory systems. Curr Opin Microbiol 61, 42–50, doi:10.1016/j.mib.2021.01.016 (2021).

29 Briegel, A. et al. Structure of bacterial cytoplasmic chemoreceptor arrays and implications for chemotactic signaling. Elife 3, e02151, doi:10.7554/eLife.02151 (2014).

30 Wuichet, K. & Zhulin, I. B. Origins and diversification of a complex signal transduction system in prokaryotes. Sci Signal 3, ra50, doi:10.1126/scisignal.2000724 (2010).

31 Stokke, R. et al. Functional interactions among filamentous Epsilonproteobacteria and Bacteroidetes in a deep-sea hydrothermal vent biofilm. Environmental microbiology 17, 4063–4077, doi:10.1111/1462-2920.12970 (2015).

32 McNally, A., Thomson, N. R., Reuter, S. & Wren, B. W. ‘Add, stir and reduce’: Yersinia spp. as model bacteria for pathogen evolution. Nature reviews. Microbiology 14, 177–190, doi:10.1038/nrmicro.2015.29 (2016).

33 Capra, E. J. & Laub, M. T. Evolution of two-component signal transduction systems. Annu Rev Microbiol 66, 325–347, doi:10.1146/annurev-micro-092611-150039 (2012).

34 Wuichet, K., Cantwell, B. J. & Zhulin, I. B. Evolution and phyletic distribution of two-component signal transduction systems. Curr Opin Microbiol 13, 219–225, doi:10.1016/j.mib.2009.12.011 (2010).

35 Ulrich, L. E. & Zhulin, I. B. The MiST2 database: a comprehensive genomics resource on microbial signal transduction. Nucleic acids research 38, D401–407, doi:10.1093/nar/gkp940 (2010).

36 Galperin, M. Y. Structural classification of bacterial response regulators: diversity of output domains and domain combinations. Journal of bacteriology 188, 4169–4182, doi:10.1128/JB.01887-05 (2006).

37 Wang, F. F. et al. A three-component signalling system fine-tunes expression kinetics of HPPK responsible for folate synthesis by positive feedback loop during stress response of Xanthomonas campestris. Environmental microbiology 16, 2126–2144, doi:10.1111/1462-2920.12293 (2014).

38 Shi, X. et al. Bioinformatics and experimental analysis of proteins of two-component systems in Myxococcus xanthus. Journal of bacteriology 190, 613–624, doi:10.1128/JB.01502-07 (2008).

39 Galperin, M. Y. Diversity of structure and function of response regulator output domains. Curr Opin Microbiol 13, 150–159, doi:10.1016/j.mib.2010.01.005 (2010).

40 Romling, U., Galperin, M. Y. & Gomelsky, M. Cyclic di-GMP: the first 25 years of a universal bacterial second messenger. Microbiology and molecular biology reviews : MMBR 77, 1–52, doi:10.1128/MMBR.00043-12 (2013).

41 Jenal, U., Reinders, A. & Lori, C. Cyclic di-GMP: second messenger extraordinaire. Nature reviews. Microbiology 15, 271–284, doi:10.1038/nrmicro.2016.190 (2017).

42 Romling, U., Gomelsky, M. & Galperin, M. Y. C-di-GMP: the dawning of a novel bacterial signalling system. Molecular microbiology 57, 629–639, doi:10.1111/j.1365-2958.2005.04697.x (2005).

43 Hengge, R. High-Specificity Local and Global c-di-GMP Signaling. Trends in microbiology, doi:10.1016/j.tim.2021.02.003 (2021).

44 Dahlstrom, K. M. & O’Toole, G. A. A Symphony of Cyclases: Specificity in Diguanylate Cyclase Signaling. Annu Rev Microbiol 71, 179–195, doi:10.1146/annurev-micro-090816-093325 (2017).

45 Hengge, R. Principles of c-di-GMP signalling in bacteria. Nature reviews. Microbiology 7, 263–273, doi:10.1038/nrmicro2109 (2009).

46 Mascher, T. Signaling diversity and evolution of extracytoplasmic function (ECF) sigma factors. Curr Opin Microbiol 16, 148–155, doi:10.1016/j.mib.2013.02.001 (2013).

47 Bassler, J., Schultz, J. E. & Lupas, A. N. Adenylate cyclases: Receivers, transducers, and generators of signals. Cell Signal 46, 135–144, doi:10.1016/j.cellsig.2018.03.002 (2018).

48 Pereira, S. F., Goss, L. & Dworkin, J. Eukaryote-like serine/threonine kinases and phosphatases in bacteria. Microbiology and molecular biology reviews : MMBR 75, 192–212, doi:10.1128/MMBR.00042-10 (2011).

49 Hickman, J. W., Tifrea, D. F. & Harwood, C. S. A chemosensory system that regulates biofilm formation through modulation of cyclic diguanylate levels. Proceedings of the National Academy of Sciences of the United States of America 102, 14422–14427, doi:10.1073/pnas.0507170102 (2005).

50 Chen, G. et al. The SiaA/B/C/D signaling network regulates biofilm formation in Pseudomonas aeruginosa. The EMBO journal 39, e103412, doi:10.15252/embj.2019103412 (2020).

51 Chen, G. et al. Structural basis for diguanylate cyclase activation by its binding partner in Pseudomonas aeruginosa. Elife 10, doi:10.7554/eLife.67289 (2021).

52 Garnerone, A. M. et al. NsrA, a Predicted beta-Barrel Outer Membrane Protein Involved in Plant Signal Perception and the Control of Secondary Infection in Sinorhizobium meliloti. Journal of bacteriology 200, doi:10.1128/JB.00019-18 (2018).

53 Capra, E. J. et al. Systematic dissection and trajectory-scanning mutagenesis of the molecular interface that ensures specificity of two-component signaling pathways. PLoS genetics 6, e1001220, doi:10.1371/journal.pgen.1001220 (2010).

54 Capra, E. J., Perchuk, B. S., Skerker, J. M. & Laub, M. T. Adaptive mutations that prevent crosstalk enable the expansion of paralogous signaling protein families. Cell 150, 222–232, doi:10.1016/j.cell.2012.05.033 (2012).

55 Farr, A. D., Remigi, P. & Rainey, P. B. Adaptive evolution by spontaneous domain fusion and protein relocalization. Nat Ecol Evol 1, 1562–1568, doi:10.1038/s41559-017-0283-7 (2017).

56 Lind, P. A., Farr, A. D. & Rainey, P. B. Experimental evolution reveals hidden diversity in evolutionary pathways. Elife 4, doi:10.7554/eLife.07074 (2015).

57 Price, M. N., Dehal, P. S. & Arkin, A. P. FastTree 2--approximately maximum-likelihood trees for large alignments. PloS one 5, e9490, doi:10.1371/journal.pone.0009490 (2010).

58 Kumar, S., Stecher, G., Li, M., Knyaz, C. & Tamura, K. MEGA X: Molecular Evolutionary Genetics Analysis across Computing Platforms. Mol Biol Evol 35, 1547–1549, doi:10.1093/molbev/msy096 (2018).

59 Camacho, C. et al. BLAST+: architecture and applications. BMC Bioinformatics 10, 421, doi:10.1186/1471-2105-10-421 (2009).

60 Rozewicki, J., Li, S., Amada, K. M., Standley, D. M. & Katoh, K. MAFFT-DASH: integrated protein sequence and structural alignment. Nucleic acids research 47, W5–W10, doi:10.1093/nar/gkz342 (2019).

61 Letunic, I., Khedkar, S. & Bork, P. SMART: recent updates, new developments and status in 2020. Nucleic acids research 49, D458–D460, doi:10.1093/nar/gkaa937 (2021).

62 Gabler, F. et al. Protein Sequence Analysis Using the MPI Bioinformatics Toolkit. Curr Protoc Bioinformatics 72, e108, doi:10.1002/cpbi.108 (2020).

63 Drinkwater, B. & Charleston, M. A. Introducing TreeCollapse: a novel greedy algorithm to solve the cophylogeny reconstruction problem. BMC Bioinformatics 15 Suppl 16, S14, doi:10.1186/1471-2105-15-S16-S14 (2014).

64 Crooks, G. E., Hon, G., Chandonia, J. M. & Brenner, S. E. WebLogo: a sequence logo generator. Genome research 14, 1188–1190, doi:10.1101/gr.849004 (2004).

65 Na, S. I. et al. UBCG: Up-to-date bacterial core gene set and pipeline for phylogenomic tree reconstruction. J Microbiol, 5, doi:10.1007/s12275-018-8014-6 (2018).

66 Hyatt, D. et al. Prodigal: prokaryotic gene recognition and translation initiation site identification. BMC Bioinformatics 11, 119, doi:10.1186/1471-2105-11-119 (2010).

67 Meng, X. & Ji, Y. Modern Computational Techniques for the HMMER Sequence Analysis. ISRN Bioinform 2013, 252183, doi:10.1155/2013/252183 (2013).

68 Talavera, G. & Castresana, J. Improvement of phylogenies after removing divergent and ambiguously aligned blocks from protein sequence alignments. Syst Biol 56, 564–577, doi:10.1080/10635150701472164 (2007).

69 Hao, Z. et al. RIdeogram: drawing SVG graphics to visualize and map genome-wide data on the idiograms. PeerJ Comput Sci 6, e251, doi:10.7717/peerj-cs.251 (2020).

70 Alexander, R. P. & Zhulin, I. B. Evolutionary genomics reveals conserved structural determinants of signaling and adaptation in microbial chemoreceptors. Proceedings of the National Academy of Sciences of the United States of America 104, 2885–2890, doi:10.1073/pnas.0609359104 (2007).

