## Supplemental Figures 1-8 for "Evolutionary Principles of Bacterial Signaling Capacity and Complexity"

**This PDF file includes:**

Extended Data Fig. 1 to 8
See separate file for Supplementary Tables 1 to 6 in one excel file

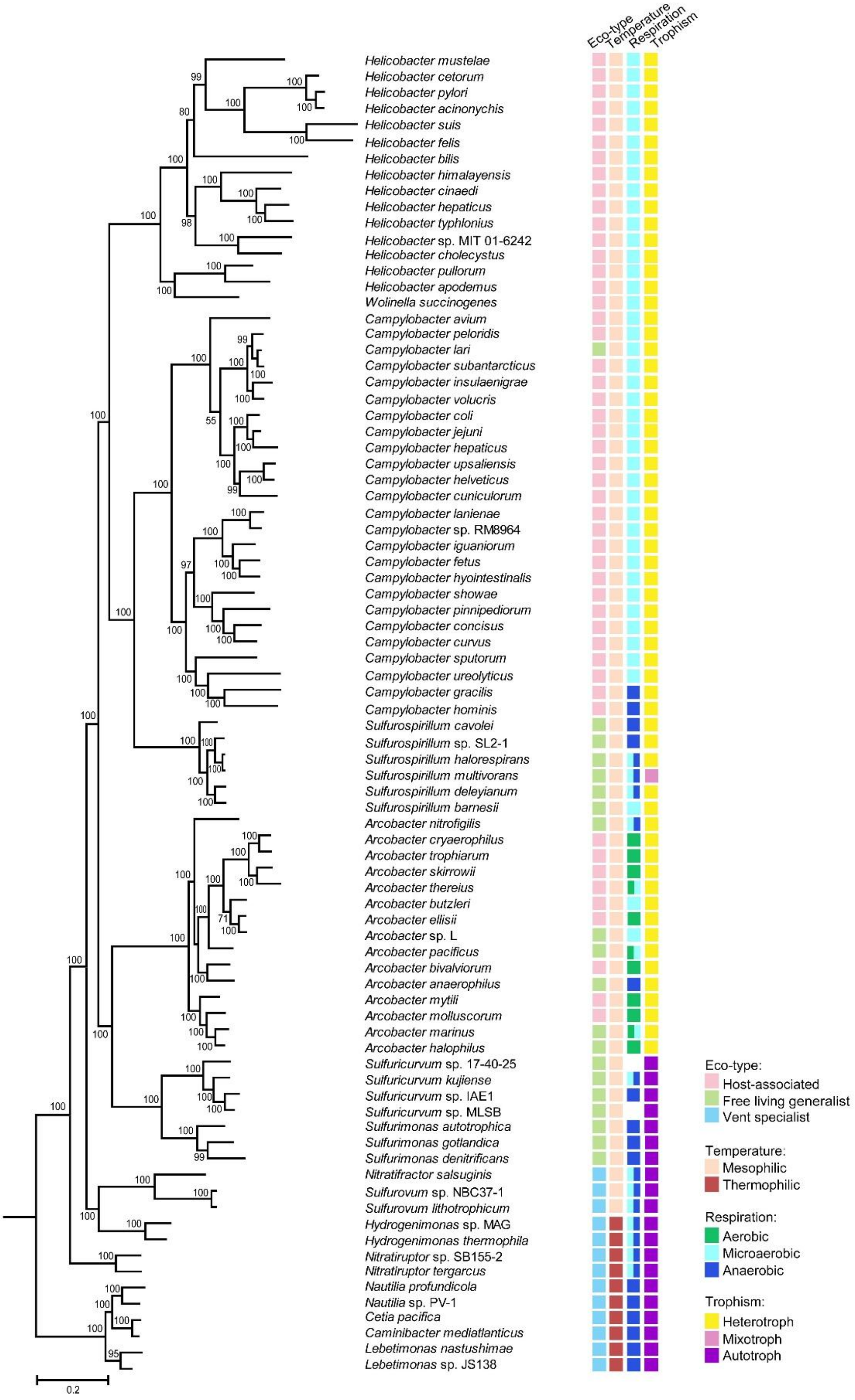


**Extended Data Fig. 1.** Phylogenetic tree and species ecophysiological characteristics of the *Campylobacterota* phylum. Detail information of all species in Supplementary Table 2.

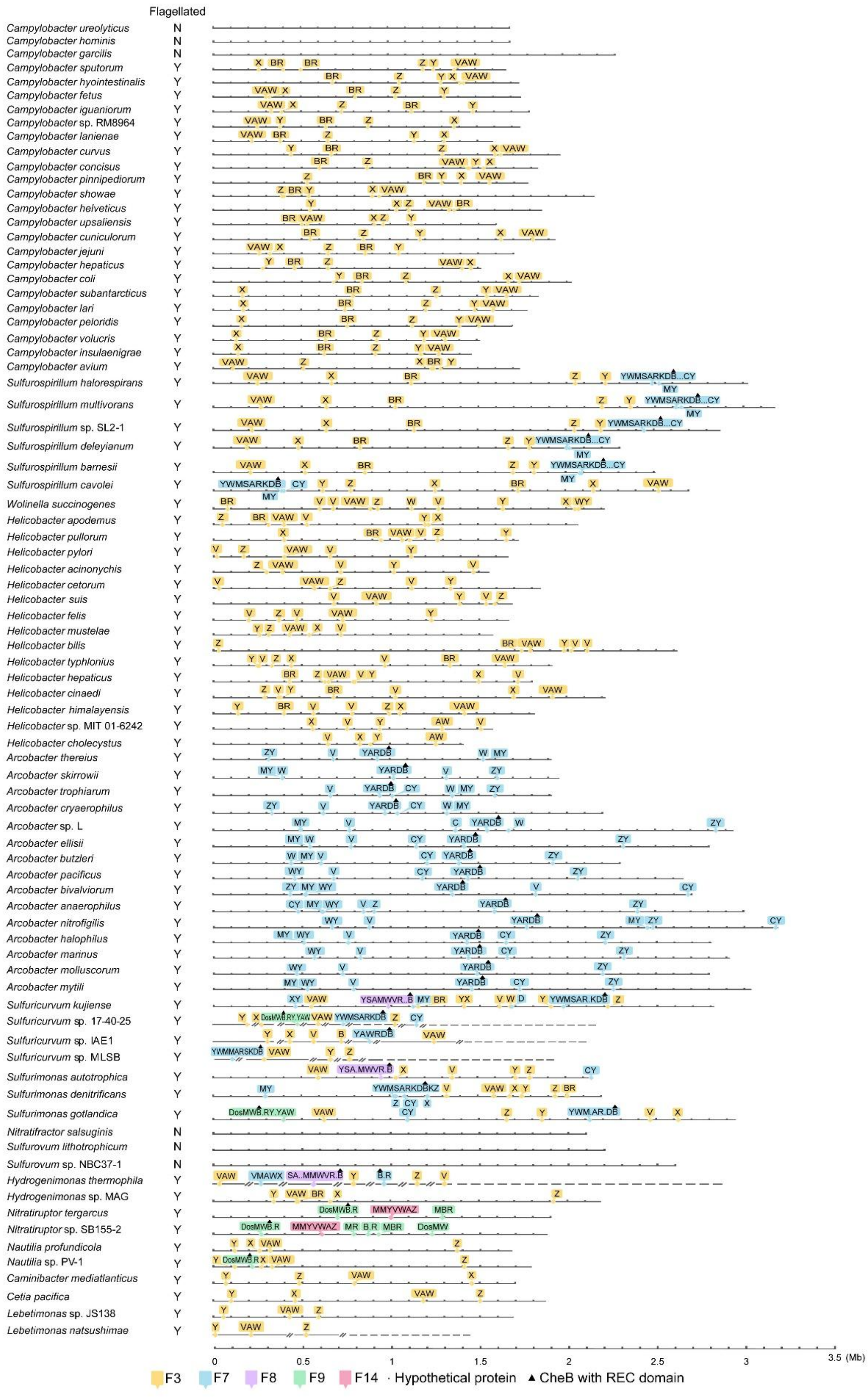

**Extended Data Fig. 2. Chemosensory classes of *Campylobacterota* species are illustrated in linearized genomes.** The color codes of chemosensory classes are noted in the bottom panel; C, *cheC*; X, *cheX*; Z, *cheZ* and other gene abbreviations are the same as Fig. 1b.

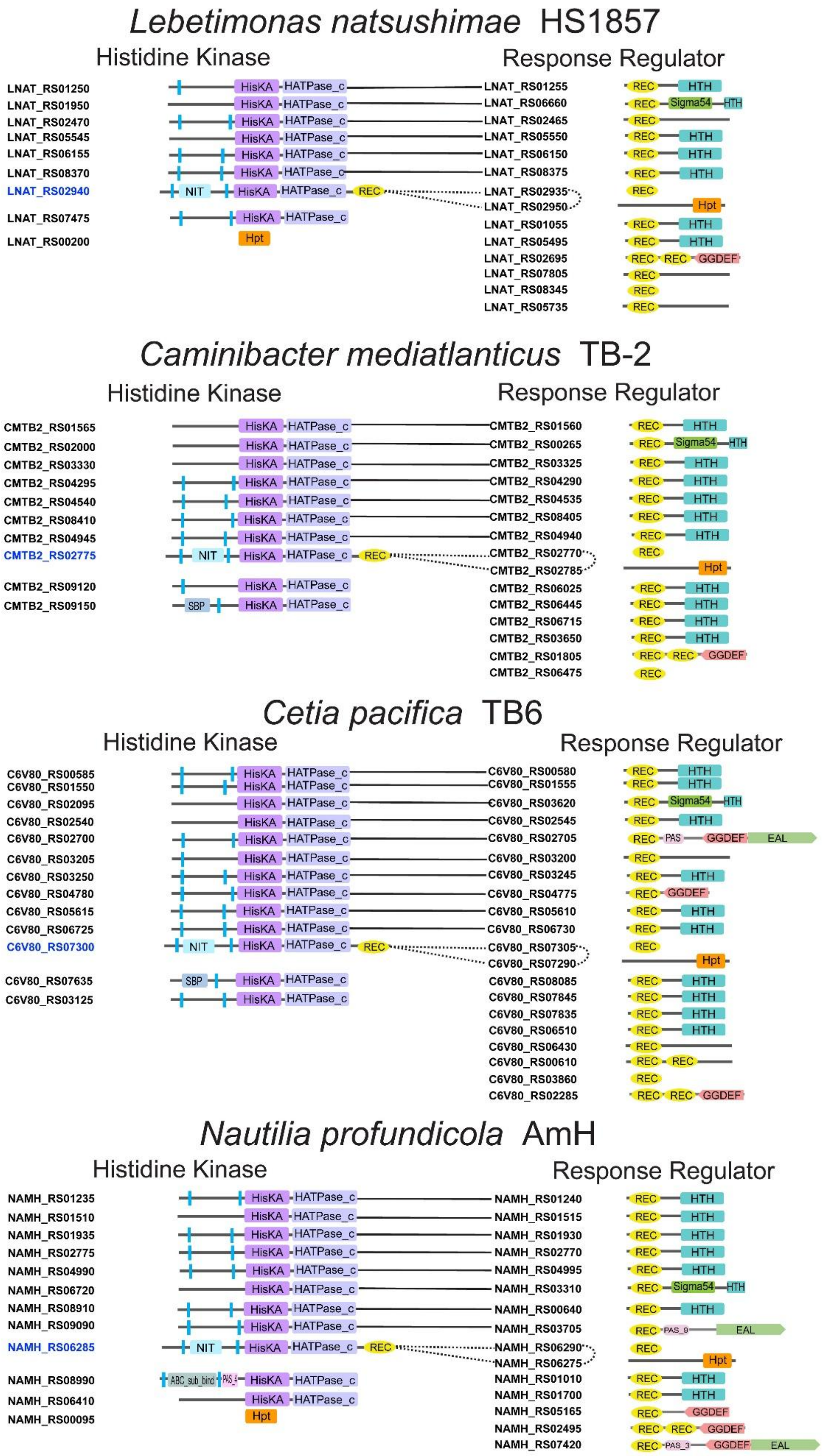
Lebetimonas natsushimae HS1857
Histidine Kinase
Response Regulator
LNAT_RS01250
LNAT_RS01950
LNAT_RS02470
LNAT_RS05545
LNAT_RS06155
LNAT_RS08370
LNAT_RS02940
LNAT_RS07475
LNAT_RS00200
LNAT_RS01255
LNAT_RS06660
LNAT_RS02465
LNAT_RS05550
LNAT_RS06150
LNAT_RS08375
LNAT_RS02935
LNAT_RS02950
LNAT_RS01055
LNAT_RS05495
LNAT_RS02695
LNAT_RS07805
LNAT_RS08345
LNAT_RS05735
Caminibacter mediatlanticus TB-2
Histidine Kinase
Response Regulator
CMTB2_RS01565
CMTB2_RS02000
CMTB2_RS03330
CMTB2_RS04295
CMTB2_RS04540
CMTB2_RS08410
CMTB2_RS04945
CMTB2_RS02775
CMTB2_RS09120
CMTB2_RS09150
CMTB2_RS01560
CMTB2_RS00265
CMTB2_RS03325
CMTB2_RS04290
CMTB2_RS04535
CMTB2_RS08405
CMTB2_RS04940
CMTB2_RS02770
CMTB2_RS02785
CMTB2_RS06025
CMTB2_RS06445
CMTB2_RS06715
CMTB2_RS03650
CMTB2_RS01805
CMTB2_RS06475
Cetia pacifica TB6
Histidine Kinase
Response Regulator
C6V80_RS00585
C6V80_RS01550
C6V80_RS02095
C6V80_RS02540
C6V80_RS02700
C6V80_RS03205
C6V80_RS03250
C6V80_RS04780
C6V80_RS05615
C6V80_RS06725
C6V80_RS07300
C6V80_RS07635
C6V80_RS03125
C6V80_RS00580
C6V80_RS01555
C6V80_RS03620
C6V80_RS02545
C6V80_RS02705
C6V80_RS03200
C6V80_RS03245
C6V80_RS04775
C6V80_RS05610
C6V80_RS06730
C6V80_RS07305
C6V80_RS07290
C6V80_RS08085
C6V80_RS07845
C6V80_RS07835
C6V80_RS06510
C6V80_RS06430
C6V80_RS00610
C6V80_RS03860
C6V80_RS02285
Nautilia profundicola AmH
Histidine Kinase
Response Regulator
NAMH_RS01235
NAMH_RS01510
NAMH_RS01935
NAMH_RS02775
NAMH_RS04990
NAMH_RS06720
NAMH_RS08910
NAMH_RS09090
NAMH_RS06285
NAMH_RS08990
NAMH_RS06410
NAMH_RS00095
NAMH_RS01240
NAMH_RS01515
NAMH_RS01930
NAMH_RS02770
NAMH_RS04995
NAMH_RS03310
NAMH_RS00640
NAMH_RS03705
NAMH_RS06290
NAMH_RS06275
NAMH_RS01010
NAMH_RS01700
NAMH_RS05165
NAMH_RS02495
NAMH_RS07420

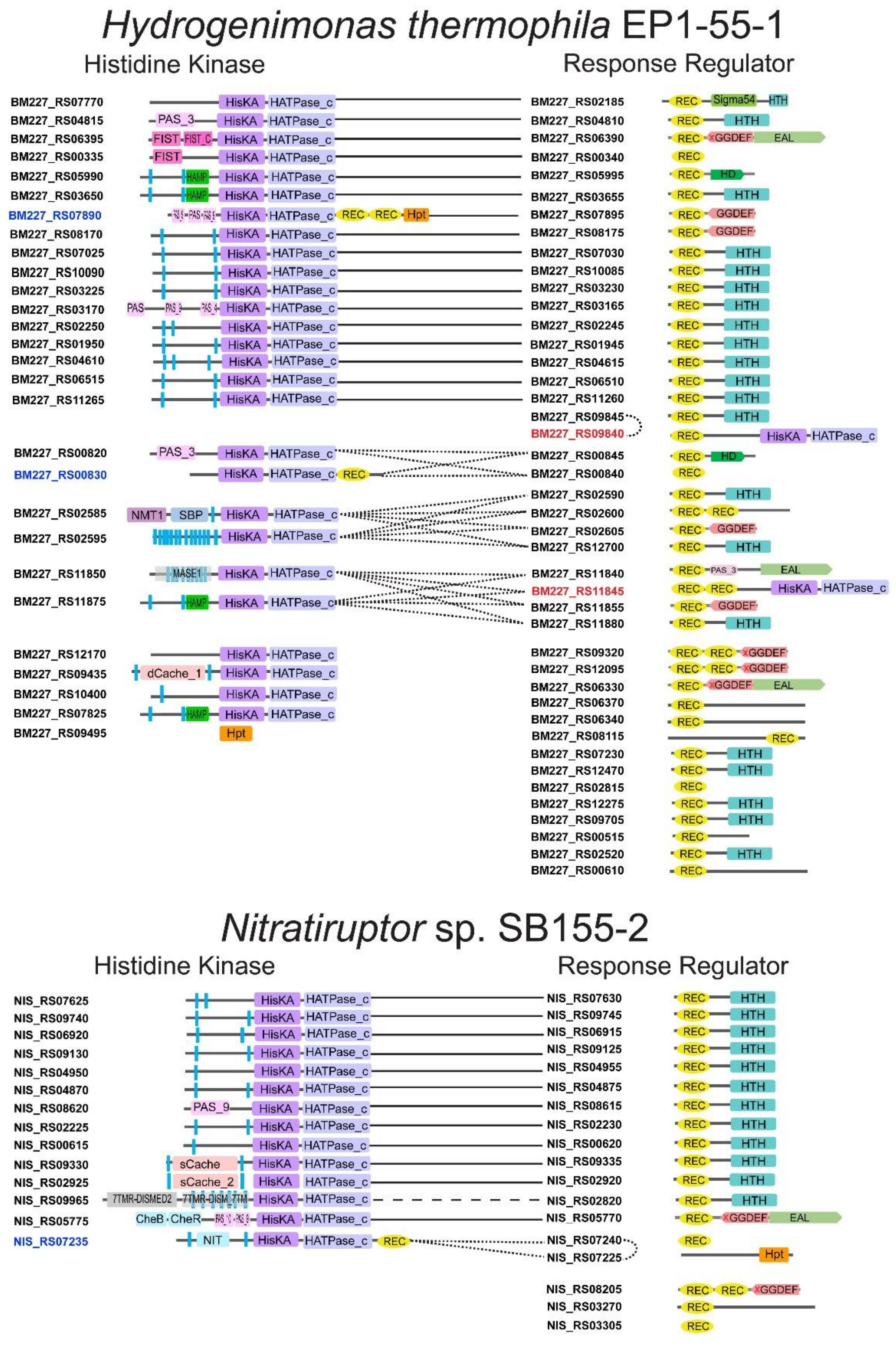

Hydrogenimonas thermophila EP1-55-1
Histidine Kinase
Response Regulator
BM227_RS07770 HisKA HATPase_c — BM227_RS02185 REC Sigma54 HTH
BM227_RS04815 PAS_3 HisKA HATPase_c — BM227_RS04810 REC HTH
BM227_RS06395 FIST FIST_C HisKA HATPase_c — BM227_RS06390 REC GGDEF EAL
BM227_RS00335 FIST HisKA HATPase_c — BM227_RS00340 REC
BM227_RS05990 HAMP HisKA HATPase_c — BM227_RS05995 REC HD
BM227_RS03650 HAMP HisKA HATPase_c — BM227_RS03655 REC HTH
BM227_RS07890 HisKA HATPase_c REC REC Hpt — BM227_RS07895 REC GGDEF
BM227_RS08170 HisKA HATPase_c — BM227_RS08175 REC GGDEF
BM227_RS07025 HisKA HATPase_c — BM227_RS07030 REC HTH
BM227_RS10090 HisKA HATPase_c — BM227_RS10085 REC HTH
BM227_RS03225 HisKA HATPase_c — BM227_RS03230 REC HTH
BM227_RS03170 PAS HisKA HATPase_c — BM227_RS03165 REC HTH
BM227_RS02250 HisKA HATPase_c — BM227_RS02245 REC HTH
BM227_RS01950 HisKA HATPase_c — BM227_RS01945 REC HTH
BM227_RS04610 HisKA HATPase_c — BM227_RS04615 REC HTH
BM227_RS06515 HisKA HATPase_c — BM227_RS06510 REC HTH
BM227_RS11265 HisKA HATPase_c — BM227_RS11260 REC HTH
BM227_RS09845 REC HTH
BM227_RS09840 REC HisKA HATPase_c
BM227_RS00820 PAS_3 HisKA HATPase_c
BM227_RS00830 HisKA HATPase_c REC
BM227_RS00845 REC HD
BM227_RS00840 REC
BM227_RS02585 NMT1 SBP HisKA HATPase_c
BM227_RS02595 HisKA HATPase_c
BM227_RS02590 REC HTH
BM227_RS02600 REC REC
BM227_RS02605 REC GGDEF
BM227_RS12700 REC HTH
BM227_RS11850 MASE1 HisKA HATPase_c
BM227_RS11875 HAMP HisKA HATPase_c
BM227_RS11840 REC PAS_3 EAL
BM227_RS11845 REC REC HisKA HATPase_c
BM227_RS11855 REC GGDEF
BM227_RS11880 REC HTH
BM227_RS12170 HisKA HATPase_c
BM227_RS09435 dCache_1 HisKA HATPase_c
BM227_RS10400 HisKA HATPase_c
BM227_RS07825 HAMP HisKA HATPase_c
BM227_RS09495 Hpt
BM227_RS09320 REC REC GGDEF
BM227_RS12095 REC REC GGDEF
BM227_RS06330 REC GGDEF EAL
BM227_RS06370 REC
BM227_RS06340 REC
BM227_RS08115 REC
BM227_RS07230 REC HTH
BM227_RS12470 REC HTH
BM227_RS02815 REC
BM227_RS12275 REC HTH
BM227_RS09705 REC HTH
BM227_RS00515 REC
BM227_RS02520 REC HTH
BM227_RS00610 REC
Nitratiruptor sp. SB155-2
Histidine Kinase
Response Regulator
NIS_RS07625 HisKA HATPase_c — NIS_RS07630 REC HTH
NIS_RS09740 HisKA HATPase_c — NIS_RS09745 REC HTH
NIS_RS06920 HisKA HATPase_c — NIS_RS06915 REC HTH
NIS_RS09130 HisKA HATPase_c — NIS_RS09125 REC HTH
NIS_RS04950 HisKA HATPase_c — NIS_RS04955 REC HTH
NIS_RS04870 HisKA HATPase_c — NIS_RS04875 REC HTH
NIS_RS08620 PAS_9 HisKA HATPase_c — NIS_RS08615 REC HTH
NIS_RS02225 HisKA HATPase_c — NIS_RS02230 REC HTH
NIS_RS00615 HisKA HATPase_c — NIS_RS00620 REC HTH
NIS_RS09330 sCache HisKA HATPase_c — NIS_RS09335 REC HTH
NIS_RS02925 sCache_2 HisKA HATPase_c — NIS_RS02920 REC HTH
NIS_RS09965 7TMR-DISMED2 7TMR-DISM_7TM HisKA HATPase_c — NIS_RS02820 REC HTH
NIS_RS05775 CheB CheR HisKA HATPase_c — NIS_RS05770 REC GGDEF EAL
NIS_RS07235 NIT HisKA HATPase_c REC
NIS_RS07240 REC
NIS_RS07225 Hpt
NIS_RS08205 REC REC GGDEF
NIS_RS03270 REC
NIS_RS03305 REC

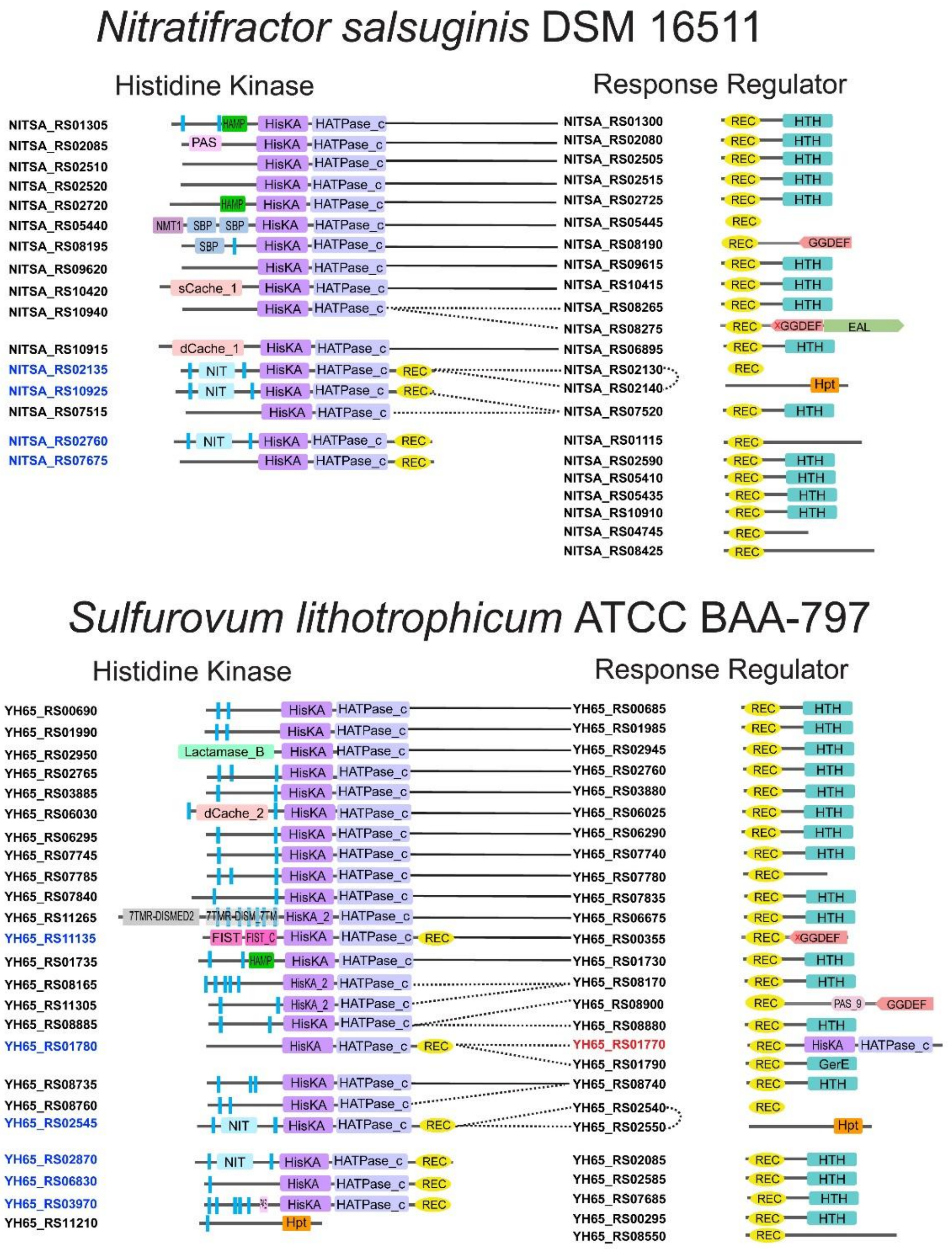

Nitratifractor salsuginis DSM 16511
Histidine Kinase
Response Regulator
NITSA_RS01305
NITSA_RS02085
NITSA_RS02510
NITSA_RS02520
NITSA_RS02720
NITSA_RS05440
NITSA_RS08195
NITSA_RS09620
NITSA_RS10420
NITSA_RS10940
NITSA_RS10915
NITSA_RS02135
NITSA_RS10925
NITSA_RS07515
NITSA_RS02760
NITSA_RS07675
NITSA_RS01300
NITSA_RS02080
NITSA_RS02505
NITSA_RS02515
NITSA_RS02725
NITSA_RS05445
NITSA_RS08190
NITSA_RS09615
NITSA_RS10415
NITSA_RS08265
NITSA_RS08275
NITSA_RS06895
NITSA_RS02130
NITSA_RS02140
NITSA_RS07520
NITSA_RS01115
NITSA_RS02590
NITSA_RS05410
NITSA_RS05435
NITSA_RS10910
NITSA_RS04745
NITSA_RS08425
HAMP
PAS
NMT1
SBP
sCache_1
dCache_1
NIT
HisKA
HATPase_c
REC
HTH
GGDEF
EAL
Hpt
Sulfurovum lithotrophicum ATCC BAA-797
Histidine Kinase
Response Regulator
YH65_RS00690
YH65_RS01990
YH65_RS02950
YH65_RS02765
YH65_RS03885
YH65_RS06030
YH65_RS06295
YH65_RS07745
YH65_RS07785
YH65_RS07840
YH65_RS11265
YH65_RS11135
YH65_RS01735
YH65_RS08165
YH65_RS11305
YH65_RS08885
YH65_RS01780
YH65_RS08735
YH65_RS08760
YH65_RS02545
YH65_RS02870
YH65_RS06830
YH65_RS03970
YH65_RS11210
YH65_RS00685
YH65_RS01985
YH65_RS02945
YH65_RS02760
YH65_RS03880
YH65_RS06025
YH65_RS06290
YH65_RS07740
YH65_RS07780
YH65_RS07835
YH65_RS06675
YH65_RS00355
YH65_RS01730
YH65_RS08170
YH65_RS08900
YH65_RS08880
YH65_RS01770
YH65_RS01790
YH65_RS08740
YH65_RS02540
YH65_RS02550
YH65_RS02085
YH65_RS02585
YH65_RS07685
YH65_RS00295
YH65_RS08550
Lactamase_B
dCache_2
7TMR-DISMED2
7TMR-DISM_7TM
FIST
FIST_C
HisKA_2
PAS_9
GerE

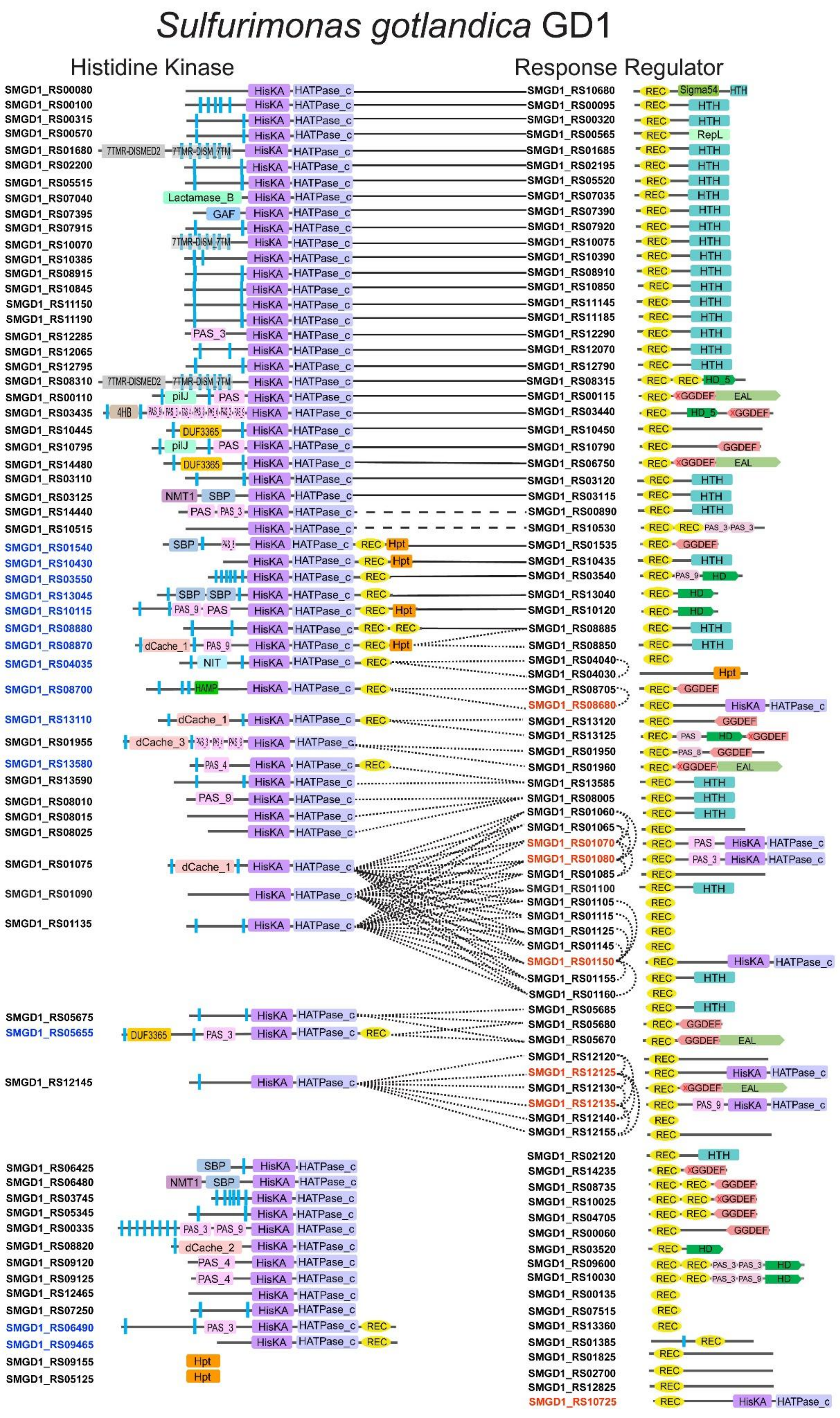

Sulfurimonas gotlandica GD1
Histidine Kinase
Response Regulator
SMGD1_RS00080
SMGD1_RS00100
SMGD1_RS00315
SMGD1_RS00570
SMGD1_RS01680
SMGD1_RS02200
SMGD1_RS05515
SMGD1_RS07040
SMGD1_RS07395
SMGD1_RS07915
SMGD1_RS10070
SMGD1_RS10385
SMGD1_RS08915
SMGD1_RS10845
SMGD1_RS11150
SMGD1_RS11190
SMGD1_RS12285
SMGD1_RS12065
SMGD1_RS12795
SMGD1_RS08310
SMGD1_RS00110
SMGD1_RS03435
SMGD1_RS10445
SMGD1_RS10795
SMGD1_RS14480
SMGD1_RS03110
SMGD1_RS03125
SMGD1_RS14440
SMGD1_RS10515
SMGD1_RS01540
SMGD1_RS10430
SMGD1_RS03550
SMGD1_RS13045
SMGD1_RS10115
SMGD1_RS08880
SMGD1_RS08870
SMGD1_RS04035
SMGD1_RS08700
SMGD1_RS13110
SMGD1_RS01955
SMGD1_RS13580
SMGD1_RS13590
SMGD1_RS08010
SMGD1_RS08015
SMGD1_RS08025
SMGD1_RS01075
SMGD1_RS01090
SMGD1_RS01135
SMGD1_RS05675
SMGD1_RS05655
SMGD1_RS12145
SMGD1_RS06425
SMGD1_RS06480
SMGD1_RS03745
SMGD1_RS05345
SMGD1_RS00335
SMGD1_RS08820
SMGD1_RS09120
SMGD1_RS09125
SMGD1_RS12465
SMGD1_RS07250
SMGD1_RS06490
SMGD1_RS09465
SMGD1_RS09155
SMGD1_RS05125
SMGD1_RS10680
SMGD1_RS00095
SMGD1_RS00320
SMGD1_RS00565
SMGD1_RS01685
SMGD1_RS02195
SMGD1_RS05520
SMGD1_RS07035
SMGD1_RS07390
SMGD1_RS07920
SMGD1_RS10075
SMGD1_RS10390
SMGD1_RS08910
SMGD1_RS10850
SMGD1_RS11145
SMGD1_RS11185
SMGD1_RS12290
SMGD1_RS12070
SMGD1_RS12790
SMGD1_RS08315
SMGD1_RS00115
SMGD1_RS03440
SMGD1_RS10450
SMGD1_RS10790
SMGD1_RS06750
SMGD1_RS03120
SMGD1_RS03115
SMGD1_RS00890
SMGD1_RS10530
SMGD1_RS01535
SMGD1_RS10435
SMGD1_RS03540
SMGD1_RS13040
SMGD1_RS10120
SMGD1_RS08885
SMGD1_RS08850
SMGD1_RS04040
SMGD1_RS04030
SMGD1_RS08705
SMGD1_RS08680
SMGD1_RS13120
SMGD1_RS13125
SMGD1_RS01950
SMGD1_RS01960
SMGD1_RS13585
SMGD1_RS08005
SMGD1_RS01060
SMGD1_RS01065
SMGD1_RS01070
SMGD1_RS01080
SMGD1_RS01085
SMGD1_RS01100
SMGD1_RS01105
SMGD1_RS01115
SMGD1_RS01125
SMGD1_RS01145
SMGD1_RS01150
SMGD1_RS01155
SMGD1_RS01160
SMGD1_RS05685
SMGD1_RS05680
SMGD1_RS05670
SMGD1_RS12120
SMGD1_RS12125
SMGD1_RS12130
SMGD1_RS12135
SMGD1_RS12140
SMGD1_RS12155
SMGD1_RS02120
SMGD1_RS14235
SMGD1_RS08735
SMGD1_RS10025
SMGD1_RS04705
SMGD1_RS00060
SMGD1_RS03520
SMGD1_RS09600
SMGD1_RS10030
SMGD1_RS00135
SMGD1_RS07515
SMGD1_RS13360
SMGD1_RS01385
SMGD1_RS01825
SMGD1_RS02700
SMGD1_RS12825
SMGD1_RS10725
HisKA
HATPase_c
REC
HTH
Hpt
Sigma54
RepL
GGDEF
EAL
HD
HD_5
PAS
PAS_3
PAS_4
PAS_8
PAS_9
GAF
SBP
NMT1
NIT
HAMP
dCache_1
dCache_2
dCache_3
DUF3365
pilJ
4HB
Lactamase_B
7TMR-DISMED2
7TMR-DISM_7TM

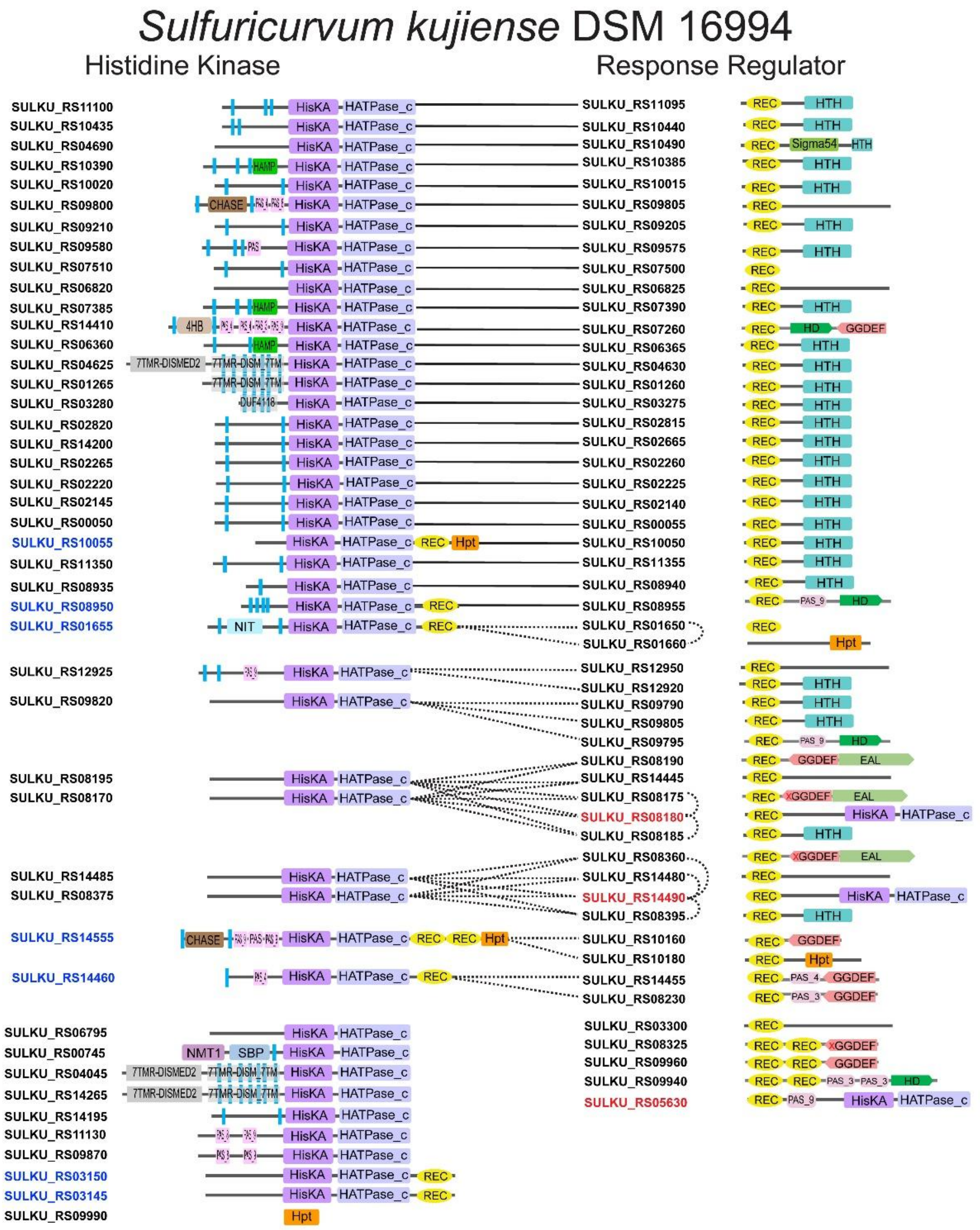

Sulfuricurvum kujiense DSM 16994
Histidine Kinase
Response Regulator
SULKU_RS11100
SULKU_RS10435
SULKU_RS04690
SULKU_RS10390
SULKU_RS10020
SULKU_RS09800
SULKU_RS09210
SULKU_RS09580
SULKU_RS07510
SULKU_RS06820
SULKU_RS07385
SULKU_RS14410
SULKU_RS06360
SULKU_RS04625
SULKU_RS01265
SULKU_RS03280
SULKU_RS02820
SULKU_RS14200
SULKU_RS02265
SULKU_RS02220
SULKU_RS02145
SULKU_RS00050
SULKU_RS10055
SULKU_RS11350
SULKU_RS08935
SULKU_RS08950
SULKU_RS01655
SULKU_RS12925
SULKU_RS09820
SULKU_RS08195
SULKU_RS08170
SULKU_RS14485
SULKU_RS08375
SULKU_RS14555
SULKU_RS14460
SULKU_RS06795
SULKU_RS00745
SULKU_RS04045
SULKU_RS14265
SULKU_RS14195
SULKU_RS11130
SULKU_RS09870
SULKU_RS03150
SULKU_RS03145
SULKU_RS09990
SULKU_RS11095
SULKU_RS10440
SULKU_RS10490
SULKU_RS10385
SULKU_RS10015
SULKU_RS09805
SULKU_RS09205
SULKU_RS09575
SULKU_RS07500
SULKU_RS06825
SULKU_RS07390
SULKU_RS07260
SULKU_RS06365
SULKU_RS04630
SULKU_RS01260
SULKU_RS03275
SULKU_RS02815
SULKU_RS02665
SULKU_RS02260
SULKU_RS02225
SULKU_RS02140
SULKU_RS00055
SULKU_RS10050
SULKU_RS11355
SULKU_RS08940
SULKU_RS08955
SULKU_RS01650
SULKU_RS01660
SULKU_RS12950
SULKU_RS12920
SULKU_RS09790
SULKU_RS09805
SULKU_RS09795
SULKU_RS08190
SULKU_RS14445
SULKU_RS08175
SULKU_RS08180
SULKU_RS08185
SULKU_RS08360
SULKU_RS14480
SULKU_RS14490
SULKU_RS08395
SULKU_RS10160
SULKU_RS10180
SULKU_RS14455
SULKU_RS08230
SULKU_RS03300
SULKU_RS08325
SULKU_RS09960
SULKU_RS09940
SULKU_RS05630
HisKA
HATPase_c
REC
HTH
Hpt
Sigma54
HAMP
CHASE
4HB
7TMR-DISMED2
7TMR-DISM_7TM
DUF4118
NIT
NMT1
SBP
PAS_9
PAS_4
PAS_3
HD
GGDEF
EAL

### *Arcobacter bivalviorum* LMG 26154

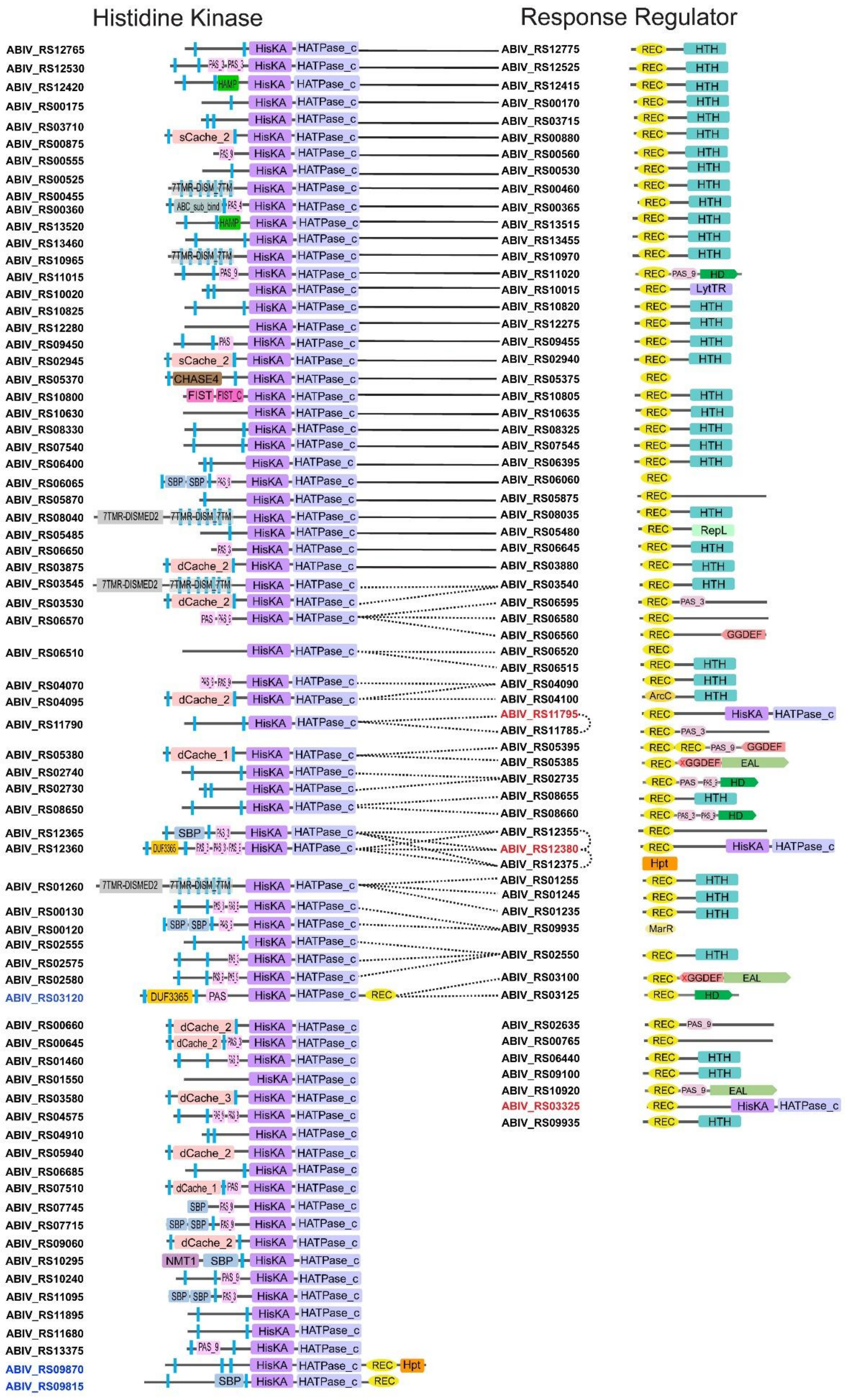

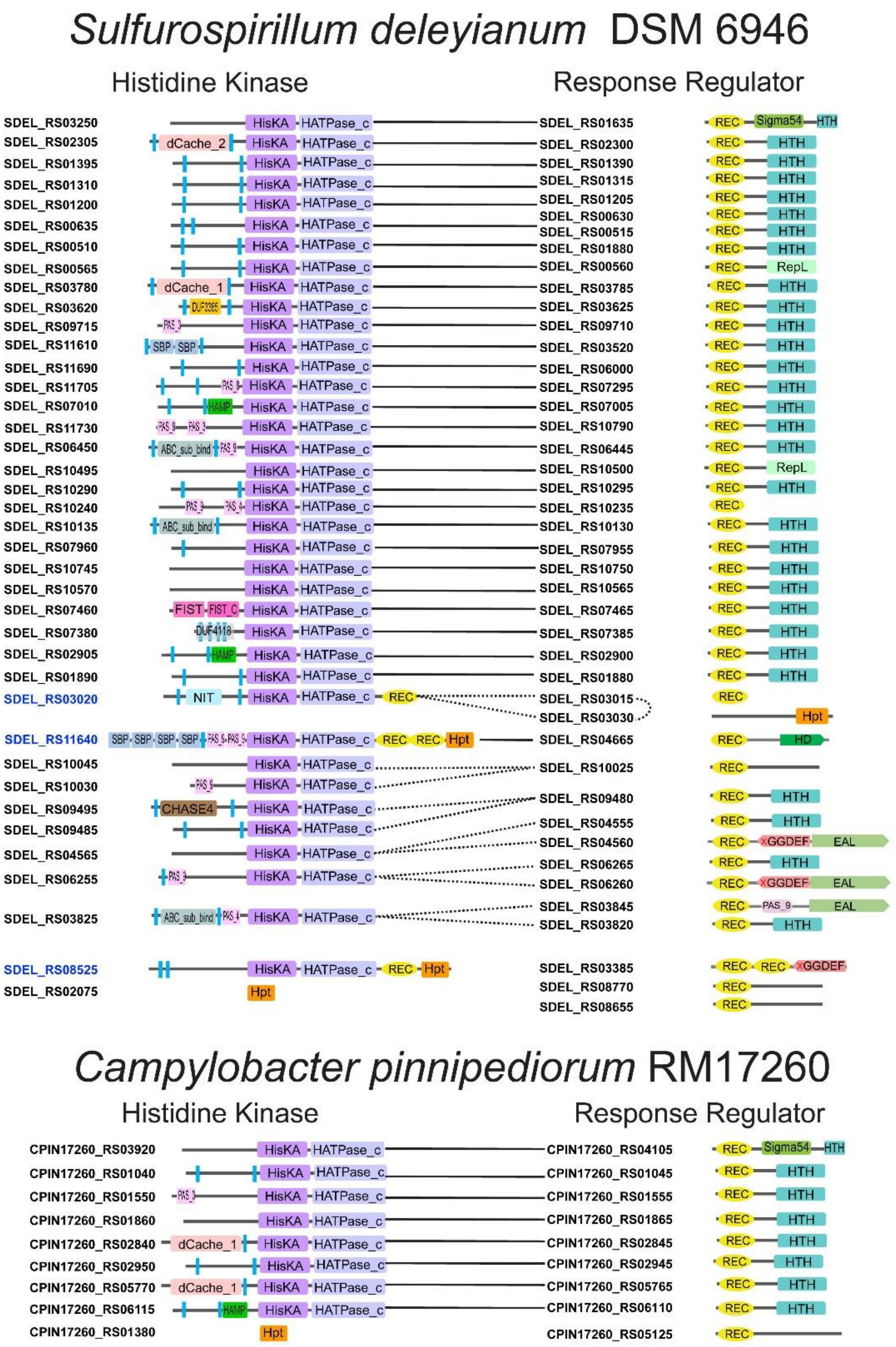

Sulfurospirillum deleyianum DSM 6946
Histidine Kinase
Response Regulator
SDEL_RS03250 HisKA HATPase_c SDEL_RS01635 REC Sigma54 HTH
SDEL_RS02305 dCache_2 HisKA HATPase_c SDEL_RS02300 REC HTH
SDEL_RS01395 HisKA HATPase_c SDEL_RS01390 REC HTH
SDEL_RS01310 HisKA HATPase_c SDEL_RS01315 REC HTH
SDEL_RS01200 HisKA HATPase_c SDEL_RS01205 REC HTH
SDEL_RS00635 HisKA HATPase_c SDEL_RS00630 REC HTH
SDEL_RS00510 HisKA HATPase_c SDEL_RS00515 REC HTH
SDEL_RS01880 REC HTH
SDEL_RS00565 HisKA HATPase_c SDEL_RS00560 REC RepL
SDEL_RS03780 dCache_1 HisKA HATPase_c SDEL_RS03785 REC HTH
SDEL_RS03620 DUF3365 HisKA HATPase_c SDEL_RS03625 REC HTH
SDEL_RS09715 PAS_3 HisKA HATPase_c SDEL_RS09710 REC HTH
SDEL_RS11610 SBP SBP HisKA HATPase_c SDEL_RS03520 REC HTH
SDEL_RS11690 HisKA HATPase_c SDEL_RS06000 REC HTH
SDEL_RS11705 PAS_9 HisKA HATPase_c SDEL_RS07295 REC HTH
SDEL_RS07010 HAMP HisKA HATPase_c SDEL_RS07005 REC HTH
SDEL_RS11730 PAS_9 PAS_3 HisKA HATPase_c SDEL_RS10790 REC HTH
SDEL_RS06450 ABC_sub_bind PAS_9 HisKA HATPase_c SDEL_RS06445 REC HTH
SDEL_RS10495 HisKA HATPase_c SDEL_RS10500 REC RepL
SDEL_RS10290 HisKA HATPase_c SDEL_RS10295 REC HTH
SDEL_RS10240 PAS_3 PAS_4 HisKA HATPase_c SDEL_RS10235 REC
SDEL_RS10135 ABC_sub_bind HisKA HATPase_c SDEL_RS10130 REC HTH
SDEL_RS07960 HisKA HATPase_c SDEL_RS07955 REC HTH
SDEL_RS10745 HisKA HATPase_c SDEL_RS10750 REC HTH
SDEL_RS10570 HisKA HATPase_c SDEL_RS10565 REC HTH
SDEL_RS07460 FIST FIST_C HisKA HATPase_c SDEL_RS07465 REC HTH
SDEL_RS07380 DUF4118 HisKA HATPase_c SDEL_RS07385 REC HTH
SDEL_RS02905 HAMP HisKA HATPase_c SDEL_RS02900 REC HTH
SDEL_RS01890 HisKA HATPase_c SDEL_RS01880 REC HTH
SDEL_RS03020 NIT HisKA HATPase_c REC SDEL_RS03015 REC
SDEL_RS03030 Hpt
SDEL_RS11640 SBP SBP SBP SBP PAS_9 PAS_9 HisKA HATPase_c REC REC Hpt SDEL_RS04665 REC HD
SDEL_RS10045 HisKA HATPase_c SDEL_RS10025 REC
SDEL_RS10030 PAS_9 HisKA HATPase_c
SDEL_RS09495 CHASE4 HisKA HATPase_c SDEL_RS09480 REC HTH
SDEL_RS09485 HisKA HATPase_c SDEL_RS04555 REC HTH
SDEL_RS04565 HisKA HATPase_c SDEL_RS04560 REC GGDEF EAL
SDEL_RS06255 PAS_3 HisKA HATPase_c SDEL_RS06265 REC HTH
SDEL_RS06260 REC GGDEF EAL
SDEL_RS03825 ABC_sub_bind PAS_4 HisKA HATPase_c SDEL_RS03845 REC PAS_9 EAL
SDEL_RS03820 REC HTH
SDEL_RS08525 HisKA HATPase_c REC Hpt SDEL_RS03385 REC REC GGDEF
SDEL_RS02075 Hpt SDEL_RS08770 REC
SDEL_RS08655 REC
Campylobacter pinnipediorum RM17260
Histidine Kinase
Response Regulator
CPIN17260_RS03920 HisKA HATPase_c CPIN17260_RS04105 REC Sigma54 HTH
CPIN17260_RS01040 HisKA HATPase_c CPIN17260_RS01045 REC HTH
CPIN17260_RS01550 PAS_3 HisKA HATPase_c CPIN17260_RS01555 REC HTH
CPIN17260_RS01860 HisKA HATPase_c CPIN17260_RS01865 REC HTH
CPIN17260_RS02840 dCache_1 HisKA HATPase_c CPIN17260_RS02845 REC HTH
CPIN17260_RS02950 HisKA HATPase_c CPIN17260_RS02945 REC HTH
CPIN17260_RS05770 dCache_1 HisKA HATPase_c CPIN17260_RS05765 REC HTH
CPIN17260_RS06115 HAMP HisKA HATPase_c CPIN17260_RS06110 REC HTH
CPIN17260_RS01380 Hpt CPIN17260_RS05125 REC

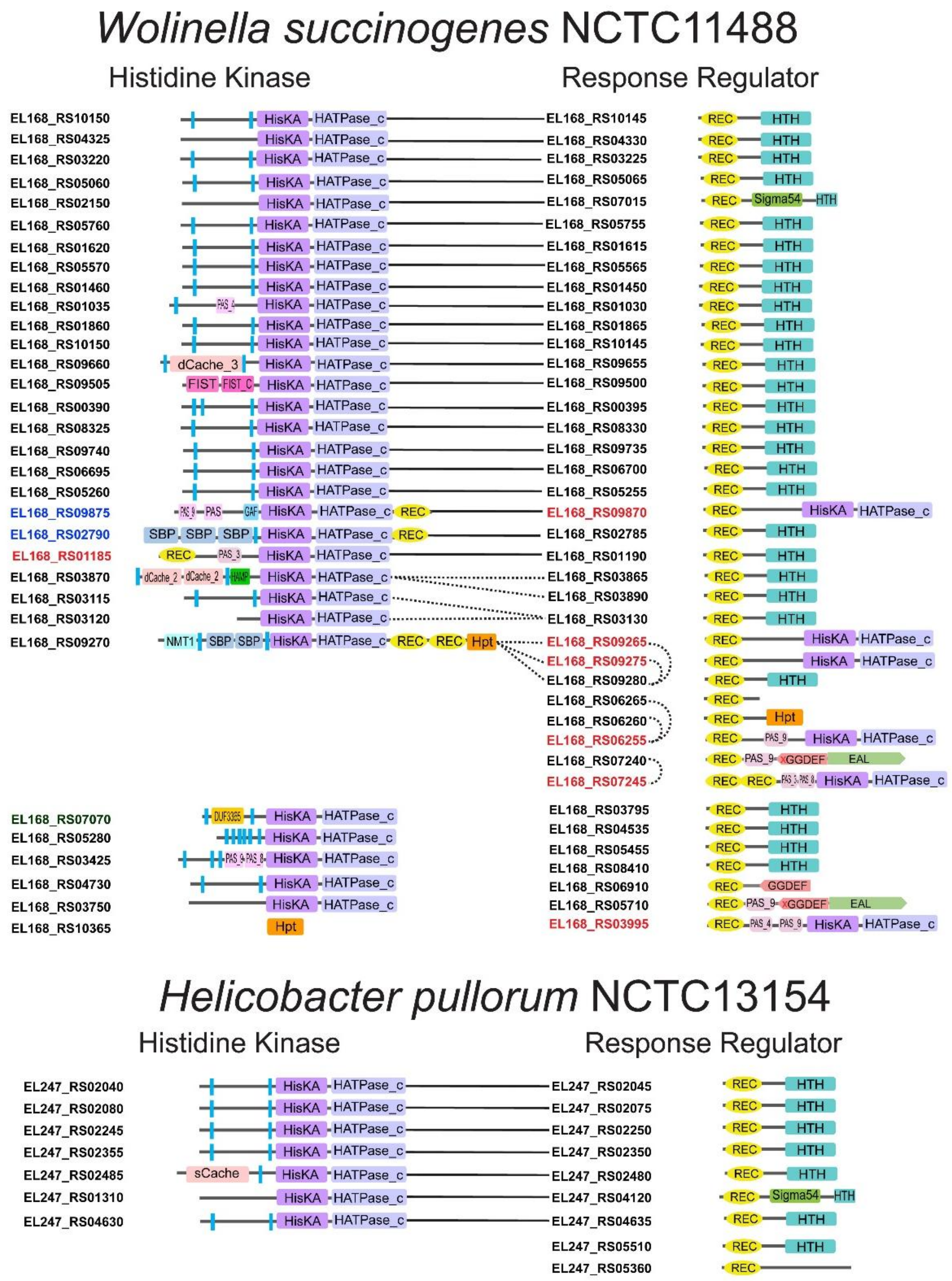


**Extended Data Fig. 3.** Domain architectures of all HKs and RRs in representative species of each genus within the *Campylobacterota* phylum. HHKs are highlighted in blue and HRRs are highlighted in red. The solid line connects the adjacent HK and RR genes; dashed line represents potential 1:1 functional link of HK and RR gene pair within four genes distance; dotted line indicates potential functional links among multiple HK and RR genes within four genes distance.

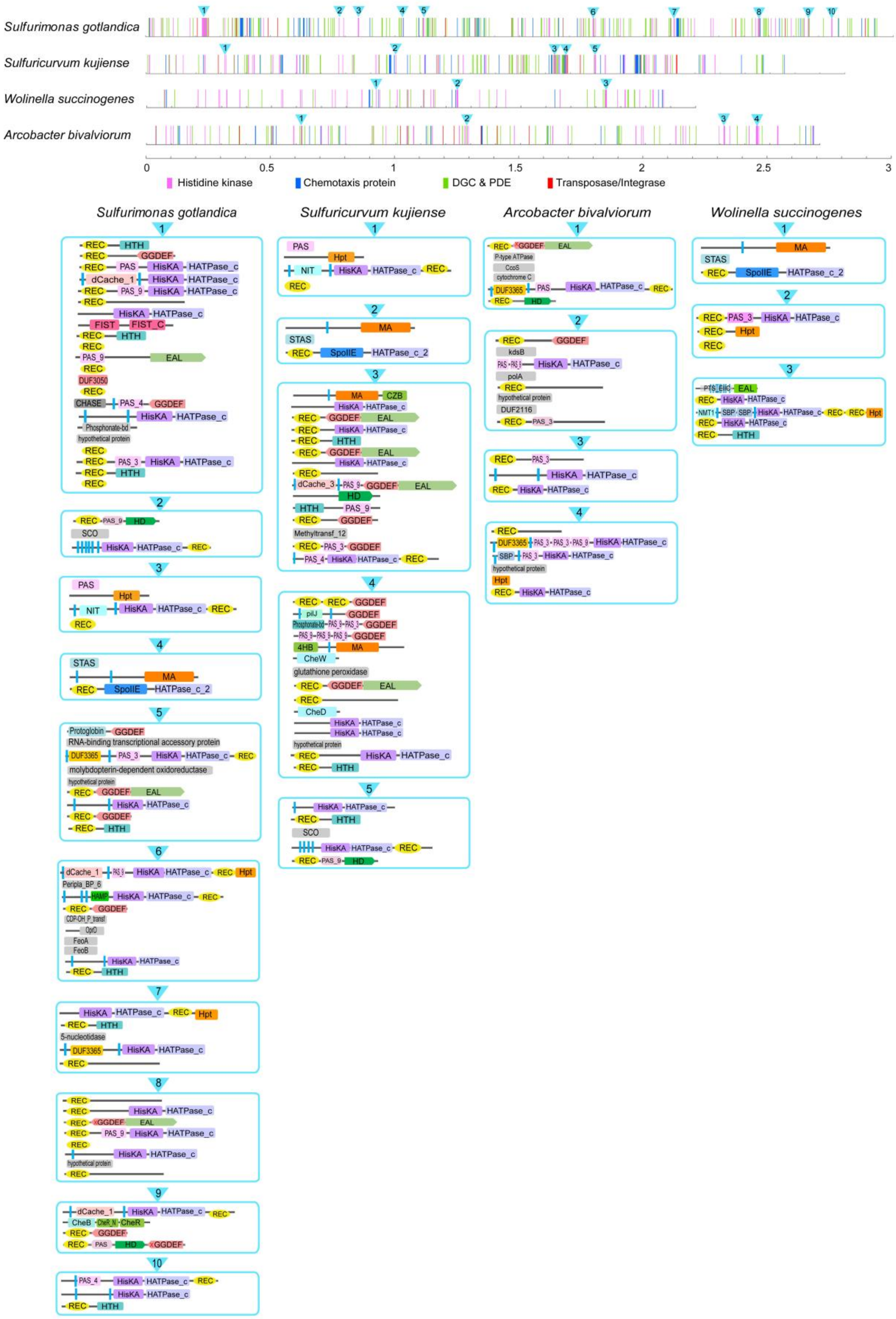


**Extended Data Fig. 4.** Atypical HK enriched clusters in four *Campylobacterota* species with abundant TCS genes. Each genome is linearized and depicted as a scale line, and signal transduction genes are represented as colored lines based on their starting location. The HHK and HRR enriched regions are illustrated at the bottom panel.

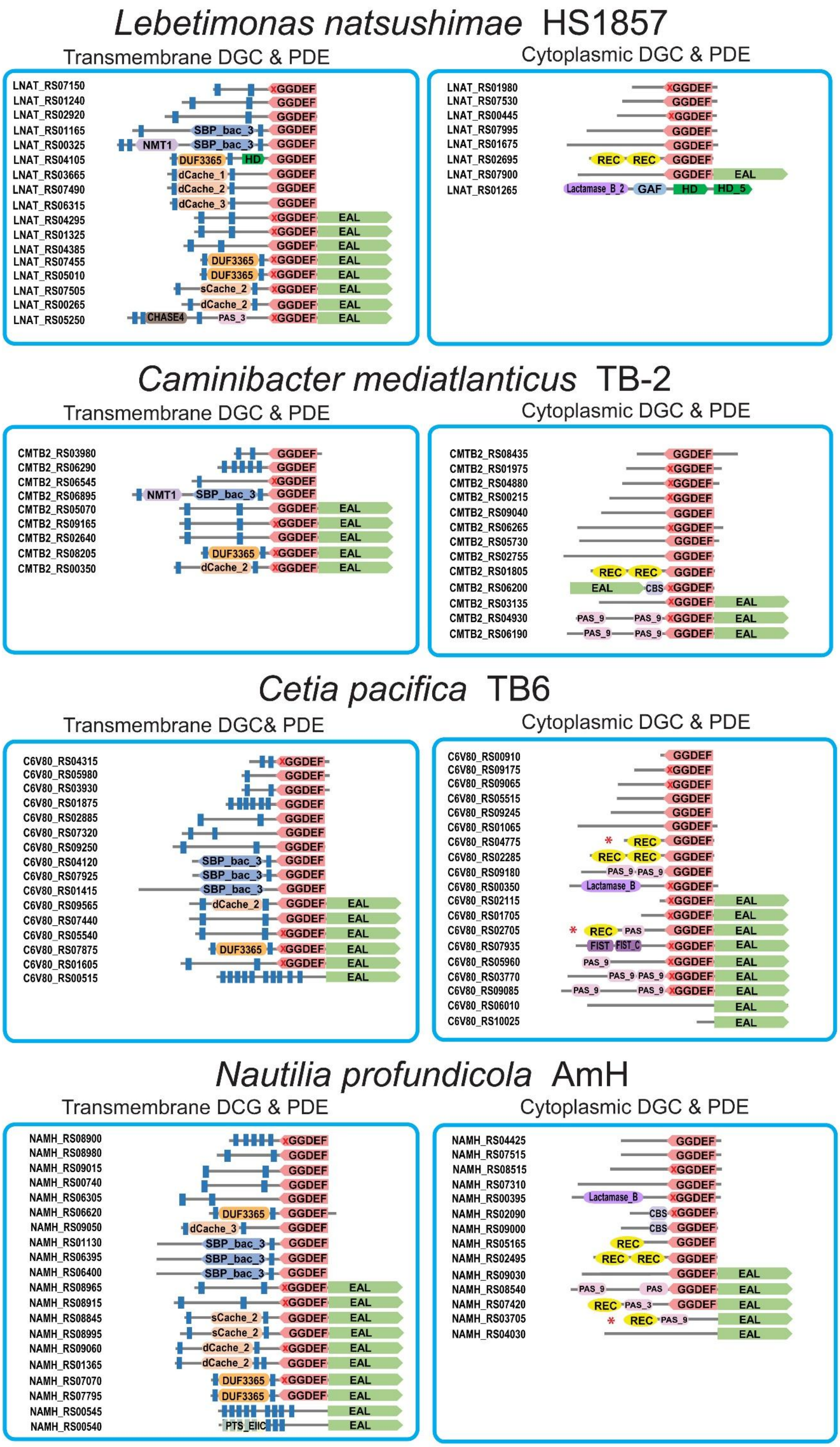

Lebetimonas natsushimae HS1857
Transmembrane DGC & PDE
LNAT_RS07150
LNAT_RS01240
LNAT_RS02920
LNAT_RS01165
LNAT_RS00325
LNAT_RS04105
LNAT_RS03665
LNAT_RS07490
LNAT_RS06315
LNAT_RS04295
LNAT_RS01325
LNAT_RS04385
LNAT_RS07455
LNAT_RS05010
LNAT_RS07505
LNAT_RS00265
LNAT_RS05250
Cytoplasmic DGC & PDE
LNAT_RS01980
LNAT_RS07530
LNAT_RS00445
LNAT_RS07995
LNAT_RS01675
LNAT_RS02695
LNAT_RS07900
LNAT_RS01265
Caminibacter mediatlanticus TB-2
Transmembrane DGC & PDE
CMTB2_RS03980
CMTB2_RS06290
CMTB2_RS06545
CMTB2_RS06895
CMTB2_RS05070
CMTB2_RS09165
CMTB2_RS02640
CMTB2_RS08205
CMTB2_RS00350
Cytoplasmic DGC & PDE
CMTB2_RS08435
CMTB2_RS01975
CMTB2_RS04880
CMTB2_RS00215
CMTB2_RS09040
CMTB2_RS06265
CMTB2_RS05730
CMTB2_RS02755
CMTB2_RS01805
CMTB2_RS06200
CMTB2_RS03135
CMTB2_RS04930
CMTB2_RS06190
Cetia pacifica TB6
Transmembrane DGC& PDE
C6V80_RS04315
C6V80_RS05980
C6V80_RS03930
C6V80_RS01875
C6V80_RS02885
C6V80_RS07320
C6V80_RS09250
C6V80_RS04120
C6V80_RS07925
C6V80_RS01415
C6V80_RS09565
C6V80_RS07440
C6V80_RS05540
C6V80_RS07875
C6V80_RS01605
C6V80_RS00515
Cytoplasmic DGC & PDE
C6V80_RS00910
C6V80_RS09175
C6V80_RS09065
C6V80_RS05515
C6V80_RS09245
C6V80_RS01065
C6V80_RS04775
C6V80_RS02285
C6V80_RS09180
C6V80_RS00350
C6V80_RS02115
C6V80_RS01705
C6V80_RS02705
C6V80_RS07935
C6V80_RS05960
C6V80_RS03770
C6V80_RS09085
C6V80_RS06010
C6V80_RS10025
Nautilia profundicola AmH
Transmembrane DCG & PDE
NAMH_RS08900
NAMH_RS08980
NAMH_RS09015
NAMH_RS00740
NAMH_RS06305
NAMH_RS06620
NAMH_RS09050
NAMH_RS01130
NAMH_RS06395
NAMH_RS06400
NAMH_RS08965
NAMH_RS08915
NAMH_RS08845
NAMH_RS08995
NAMH_RS09060
NAMH_RS01365
NAMH_RS07070
NAMH_RS07795
NAMH_RS00545
NAMH_RS00540
Cytoplasmic DGC & PDE
NAMH_RS04425
NAMH_RS07515
NAMH_RS08515
NAMH_RS07310
NAMH_RS00395
NAMH_RS02090
NAMH_RS09000
NAMH_RS05165
NAMH_RS02495
NAMH_RS09030
NAMH_RS08540
NAMH_RS07420
NAMH_RS03705
NAMH_RS04030
GGDEF
EAL
REC
PAS_9
SBP_bac_3
DUF3365
dCache_2

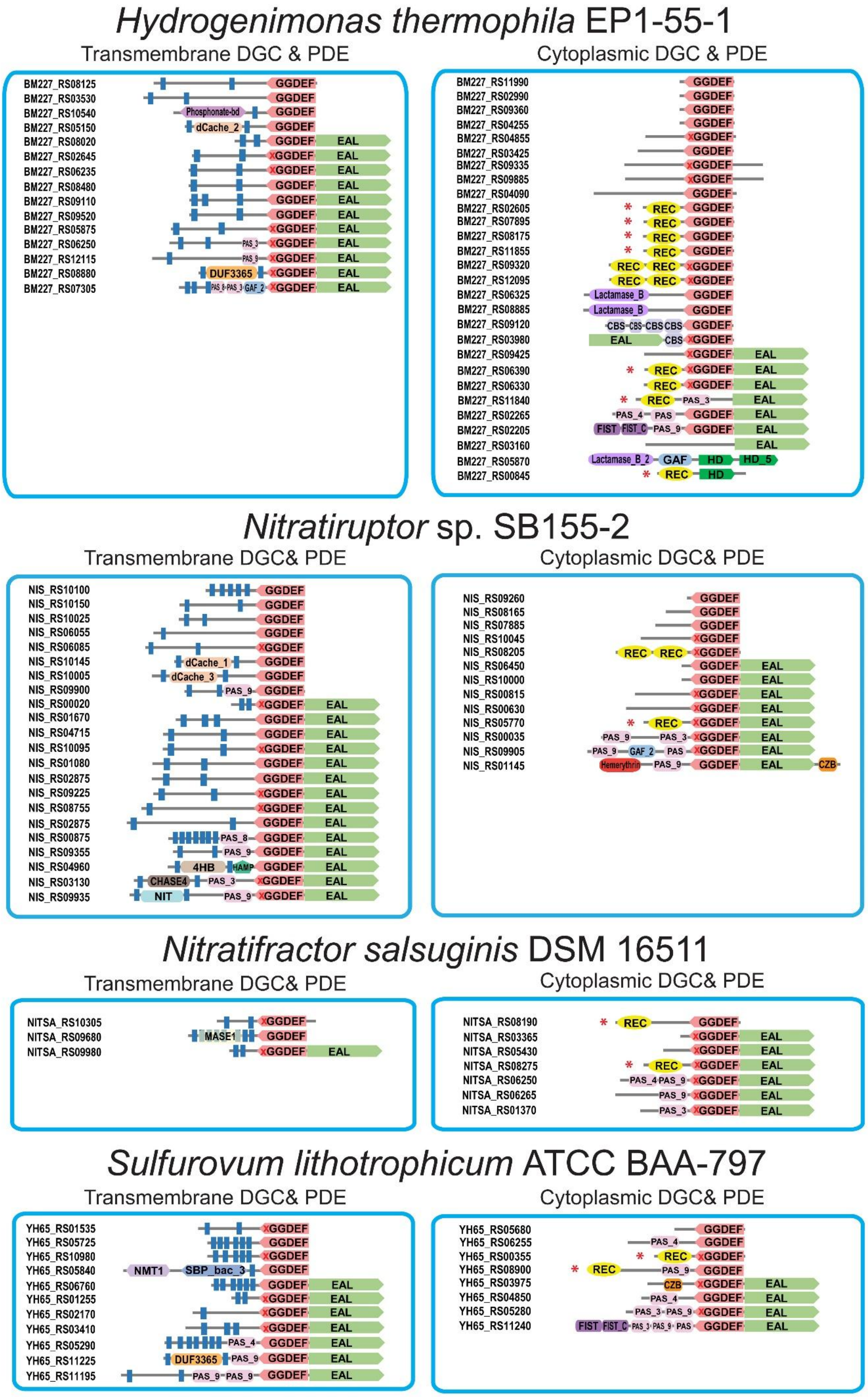

Hydrogenimonas thermophila EP1-55-1
Transmembrane DGC & PDE
BM227_RS08125
BM227_RS03530
BM227_RS10540
BM227_RS05150
BM227_RS08020
BM227_RS02645
BM227_RS06235
BM227_RS08480
BM227_RS09110
BM227_RS09520
BM227_RS05875
BM227_RS06250
BM227_RS12115
BM227_RS08880
BM227_RS07305
Cytoplasmic DGC & PDE
BM227_RS11990
BM227_RS02990
BM227_RS09360
BM227_RS04255
BM227_RS04855
BM227_RS03425
BM227_RS09335
BM227_RS09885
BM227_RS04090
BM227_RS02605
BM227_RS07895
BM227_RS08175
BM227_RS11855
BM227_RS09320
BM227_RS12095
BM227_RS06325
BM227_RS08885
BM227_RS09120
BM227_RS03980
BM227_RS09425
BM227_RS06390
BM227_RS06330
BM227_RS11840
BM227_RS02265
BM227_RS02205
BM227_RS03160
BM227_RS05870
BM227_RS00845
Nitratiruptor sp. SB155-2
Transmembrane DGC& PDE
NIS_RS10100
NIS_RS10150
NIS_RS10025
NIS_RS06055
NIS_RS06085
NIS_RS10145
NIS_RS10005
NIS_RS09900
NIS_RS00020
NIS_RS01670
NIS_RS04715
NIS_RS10095
NIS_RS01080
NIS_RS02875
NIS_RS09225
NIS_RS08755
NIS_RS02875
NIS_RS00875
NIS_RS09355
NIS_RS04960
NIS_RS03130
NIS_RS09935
Cytoplasmic DGC& PDE
NIS_RS09260
NIS_RS08165
NIS_RS07885
NIS_RS10045
NIS_RS08205
NIS_RS06450
NIS_RS10000
NIS_RS00815
NIS_RS00630
NIS_RS05770
NIS_RS00035
NIS_RS09905
NIS_RS01145
Nitratifractor salsuginis DSM 16511
Transmembrane DGC& PDE
NITSA_RS10305
NITSA_RS09680
NITSA_RS09980
Cytoplasmic DGC& PDE
NITSA_RS08190
NITSA_RS03365
NITSA_RS05430
NITSA_RS08275
NITSA_RS06250
NITSA_RS06265
NITSA_RS01370
Sulfurovum lithotrophicum ATCC BAA-797
Transmembrane DGC& PDE
YH65_RS01535
YH65_RS05725
YH65_RS10980
YH65_RS05840
YH65_RS06760
YH65_RS01255
YH65_RS02170
YH65_RS03410
YH65_RS05290
YH65_RS11225
YH65_RS11195
Cytoplasmic DGC& PDE
YH65_RS05680
YH65_RS06255
YH65_RS00355
YH65_RS08900
YH65_RS03975
YH65_RS04850
YH65_RS05280
YH65_RS11240
GGDEF
EAL
REC
PAS_3
PAS_4
PAS_8
PAS_9
GAF_2
DUF3365
dCache_1
dCache_2
dCache_3
Phosphonate-bd
Lactamase_B
Lactamase_B_2
CBS
FIST
FIST_C
GAF
HD
HD_5
Hemerythrin
CZB
4HB
HAMP
CHASE4
NIT
MASE1
NMT1
SBP_bac_3

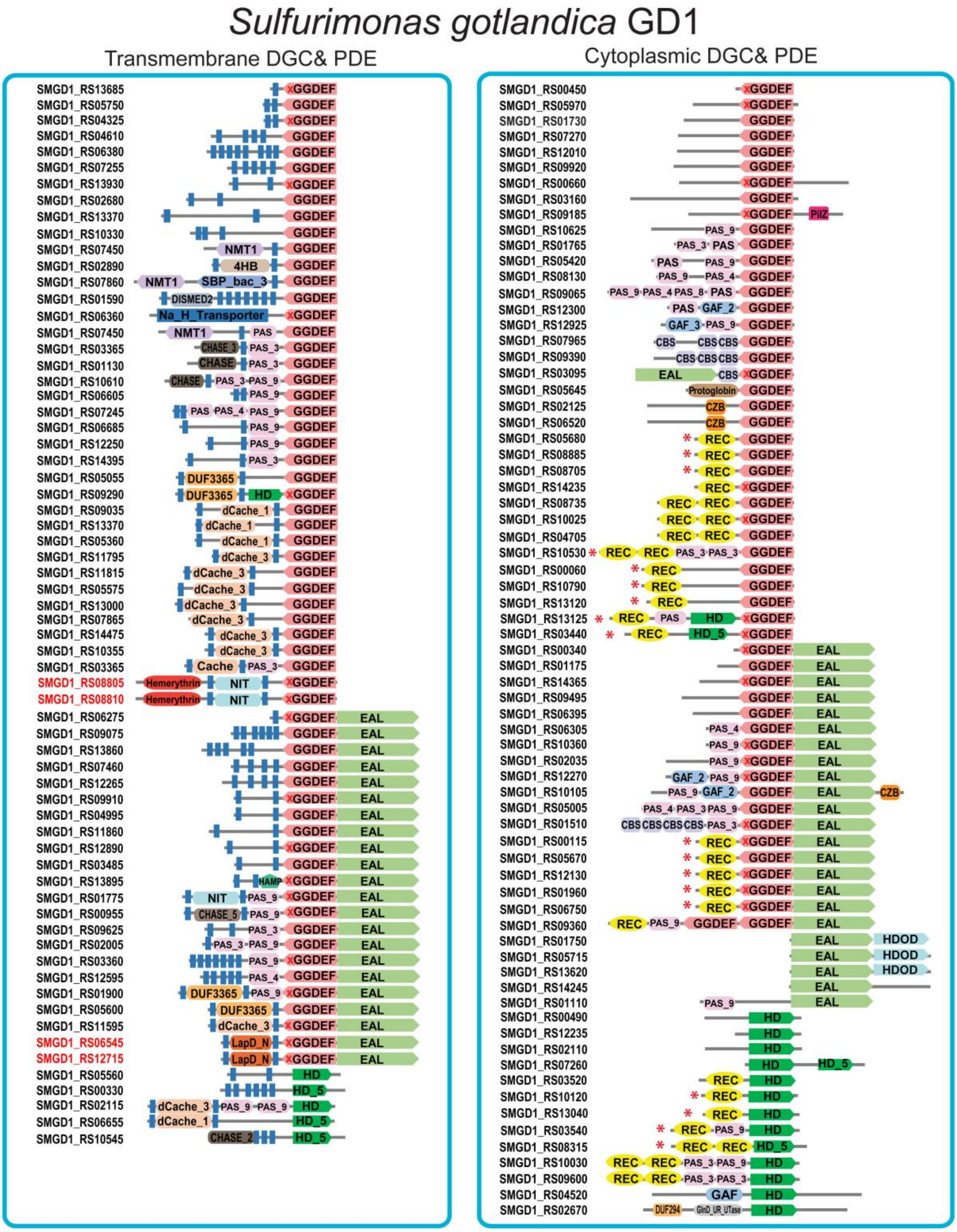

Sulfurimonas gotlandica GD1
Transmembrane DGC& PDE
Cytoplasmic DGC& PDE
SMGD1_RS13685
SMGD1_RS05750
SMGD1_RS04325
SMGD1_RS04610
SMGD1_RS06380
SMGD1_RS07255
SMGD1_RS13930
SMGD1_RS02680
SMGD1_RS13370
SMGD1_RS10330
SMGD1_RS07450
SMGD1_RS02890
SMGD1_RS07860
SMGD1_RS01590
SMGD1_RS06360
SMGD1_RS07450
SMGD1_RS03365
SMGD1_RS01130
SMGD1_RS10610
SMGD1_RS06605
SMGD1_RS07245
SMGD1_RS06685
SMGD1_RS12250
SMGD1_RS14395
SMGD1_RS05055
SMGD1_RS09290
SMGD1_RS09035
SMGD1_RS13370
SMGD1_RS05360
SMGD1_RS11795
SMGD1_RS11815
SMGD1_RS05575
SMGD1_RS13000
SMGD1_RS07865
SMGD1_RS14475
SMGD1_RS10355
SMGD1_RS03365
SMGD1_RS08805
SMGD1_RS08810
SMGD1_RS06275
SMGD1_RS09075
SMGD1_RS13860
SMGD1_RS07460
SMGD1_RS12265
SMGD1_RS09910
SMGD1_RS04995
SMGD1_RS11860
SMGD1_RS12890
SMGD1_RS03485
SMGD1_RS13895
SMGD1_RS01775
SMGD1_RS00955
SMGD1_RS09625
SMGD1_RS02005
SMGD1_RS03360
SMGD1_RS12595
SMGD1_RS01900
SMGD1_RS05600
SMGD1_RS11595
SMGD1_RS06545
SMGD1_RS12715
SMGD1_RS05560
SMGD1_RS00330
SMGD1_RS02115
SMGD1_RS06655
SMGD1_RS10545
SMGD1_RS00450
SMGD1_RS05970
SMGD1_RS01730
SMGD1_RS07270
SMGD1_RS12010
SMGD1_RS09920
SMGD1_RS00660
SMGD1_RS03160
SMGD1_RS09185
SMGD1_RS10625
SMGD1_RS01765
SMGD1_RS05420
SMGD1_RS08130
SMGD1_RS09065
SMGD1_RS12300
SMGD1_RS12925
SMGD1_RS07965
SMGD1_RS09390
SMGD1_RS03095
SMGD1_RS05645
SMGD1_RS02125
SMGD1_RS06520
SMGD1_RS05680
SMGD1_RS08885
SMGD1_RS08705
SMGD1_RS14235
SMGD1_RS08735
SMGD1_RS10025
SMGD1_RS04705
SMGD1_RS10530
SMGD1_RS00060
SMGD1_RS10790
SMGD1_RS13120
SMGD1_RS13125
SMGD1_RS03440
SMGD1_RS00340
SMGD1_RS01175
SMGD1_RS14365
SMGD1_RS09495
SMGD1_RS06395
SMGD1_RS06305
SMGD1_RS10360
SMGD1_RS02035
SMGD1_RS12270
SMGD1_RS10105
SMGD1_RS05005
SMGD1_RS01510
SMGD1_RS00115
SMGD1_RS05670
SMGD1_RS12130
SMGD1_RS01960
SMGD1_RS06750
SMGD1_RS09360
SMGD1_RS01750
SMGD1_RS05715
SMGD1_RS13620
SMGD1_RS14245
SMGD1_RS01110
SMGD1_RS00490
SMGD1_RS12235
SMGD1_RS02110
SMGD1_RS07260
SMGD1_RS03520
SMGD1_RS10120
SMGD1_RS13040
SMGD1_RS03540
SMGD1_RS08315
SMGD1_RS10030
SMGD1_RS09600
SMGD1_RS04520
SMGD1_RS02670

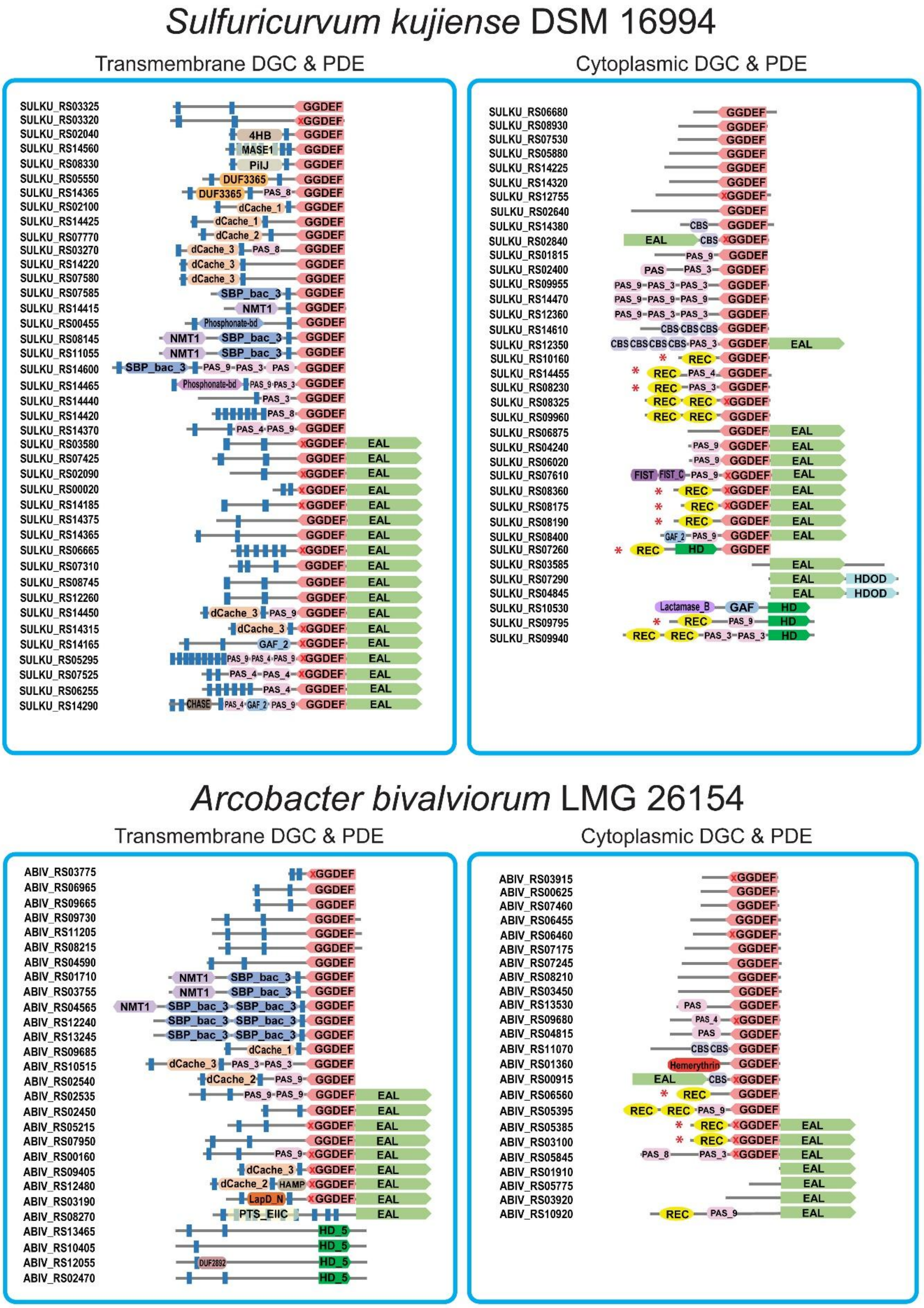
Sulfuricurvum kujiense DSM 16994
Transmembrane DGC & PDE
Cytoplasmic DGC & PDE
SULKU_RS03325
SULKU_RS03320
SULKU_RS02040
SULKU_RS14560
SULKU_RS08330
SULKU_RS05550
SULKU_RS14365
SULKU_RS02100
SULKU_RS14425
SULKU_RS07770
SULKU_RS03270
SULKU_RS14220
SULKU_RS07580
SULKU_RS07585
SULKU_RS14415
SULKU_RS00455
SULKU_RS08145
SULKU_RS11055
SULKU_RS14600
SULKU_RS14465
SULKU_RS14440
SULKU_RS14420
SULKU_RS14370
SULKU_RS03580
SULKU_RS07425
SULKU_RS02090
SULKU_RS00020
SULKU_RS14185
SULKU_RS14375
SULKU_RS14365
SULKU_RS06665
SULKU_RS07310
SULKU_RS08745
SULKU_RS12260
SULKU_RS14450
SULKU_RS14315
SULKU_RS14165
SULKU_RS05295
SULKU_RS07525
SULKU_RS06255
SULKU_RS14290
SULKU_RS06680
SULKU_RS08930
SULKU_RS07530
SULKU_RS05880
SULKU_RS14225
SULKU_RS14320
SULKU_RS12755
SULKU_RS02640
SULKU_RS14380
SULKU_RS02840
SULKU_RS01815
SULKU_RS02400
SULKU_RS09955
SULKU_RS14470
SULKU_RS12360
SULKU_RS14610
SULKU_RS12350
SULKU_RS10160
SULKU_RS14455
SULKU_RS08230
SULKU_RS08325
SULKU_RS09960
SULKU_RS06875
SULKU_RS04240
SULKU_RS06020
SULKU_RS07610
SULKU_RS08360
SULKU_RS08175
SULKU_RS08190
SULKU_RS08400
SULKU_RS07260
SULKU_RS03585
SULKU_RS07290
SULKU_RS04845
SULKU_RS10530
SULKU_RS09795
SULKU_RS09940
Arcobacter bivalviorum LMG 26154
Transmembrane DGC & PDE
Cytoplasmic DGC & PDE
ABIV_RS03775
ABIV_RS06965
ABIV_RS09665
ABIV_RS09730
ABIV_RS11205
ABIV_RS08215
ABIV_RS04590
ABIV_RS01710
ABIV_RS03755
ABIV_RS04565
ABIV_RS12240
ABIV_RS13245
ABIV_RS09685
ABIV_RS10515
ABIV_RS02540
ABIV_RS02535
ABIV_RS02450
ABIV_RS05215
ABIV_RS07950
ABIV_RS00160
ABIV_RS09405
ABIV_RS12480
ABIV_RS03190
ABIV_RS08270
ABIV_RS13465
ABIV_RS10405
ABIV_RS12055
ABIV_RS02470
ABIV_RS03915
ABIV_RS00625
ABIV_RS07460
ABIV_RS06455
ABIV_RS06460
ABIV_RS07175
ABIV_RS07245
ABIV_RS08210
ABIV_RS03450
ABIV_RS13530
ABIV_RS09680
ABIV_RS04815
ABIV_RS11070
ABIV_RS01360
ABIV_RS00915
ABIV_RS06560
ABIV_RS05395
ABIV_RS05385
ABIV_RS03100
ABIV_RS05845
ABIV_RS01910
ABIV_RS05775
ABIV_RS03920
ABIV_RS10920

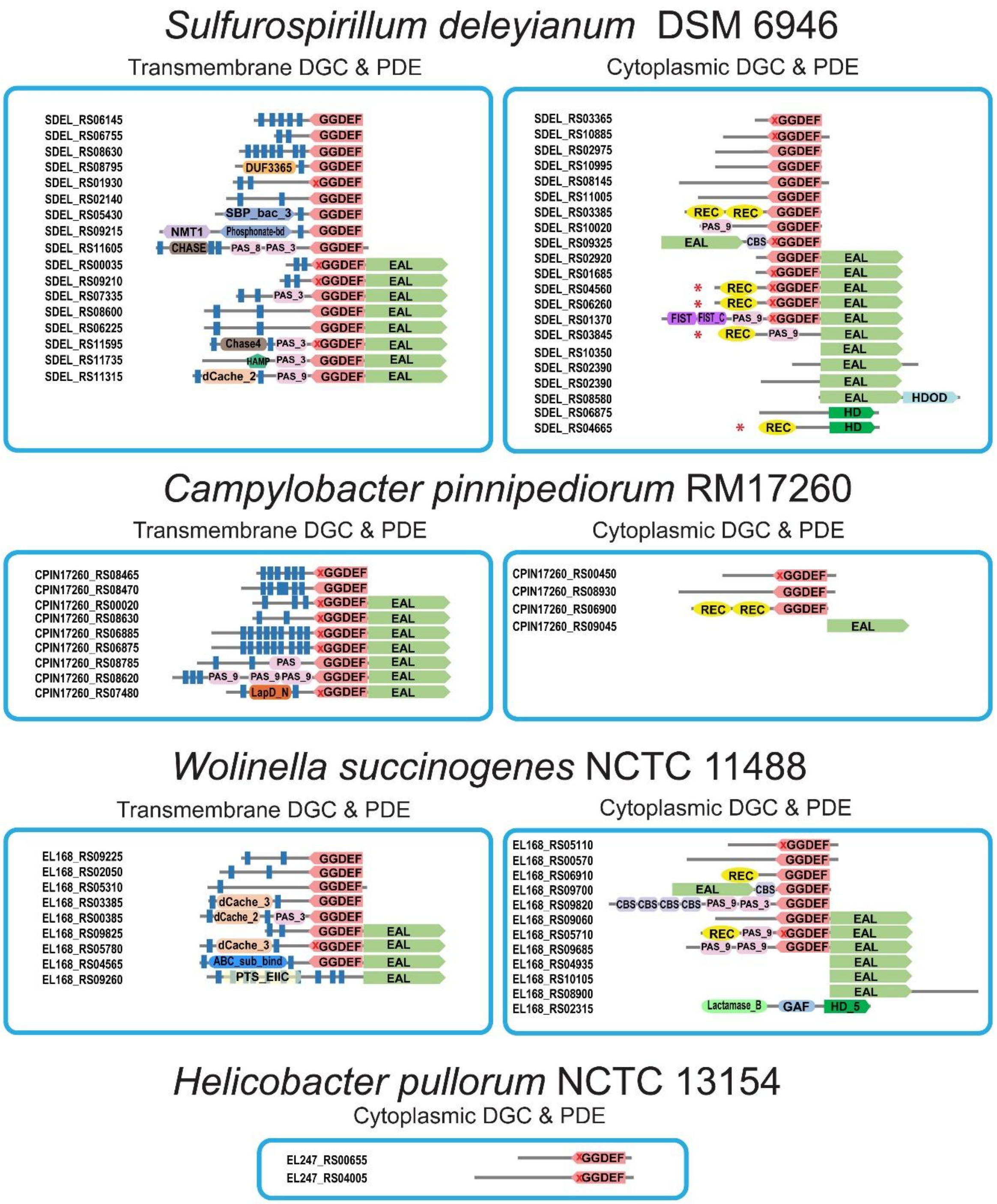


**Extended Data Fig. 5.** Domain architectures of all DGCs and PDEs in representative species of each genus within the *Campylobacterota* phylum. These proteins are categorized as transmembrane and cytoplasmic proteins. The red asterisk represents a DGC or PDE gene with a REC domain, also in close proximity of a HK gene. The red cross in the GGDEF domain implies an enzymatically inactive DGC. In the box of *Sulfurimonas gotlandica* GD1, two pairs of DGCs are highlighted in red indicating recent gene duplication of two genes.

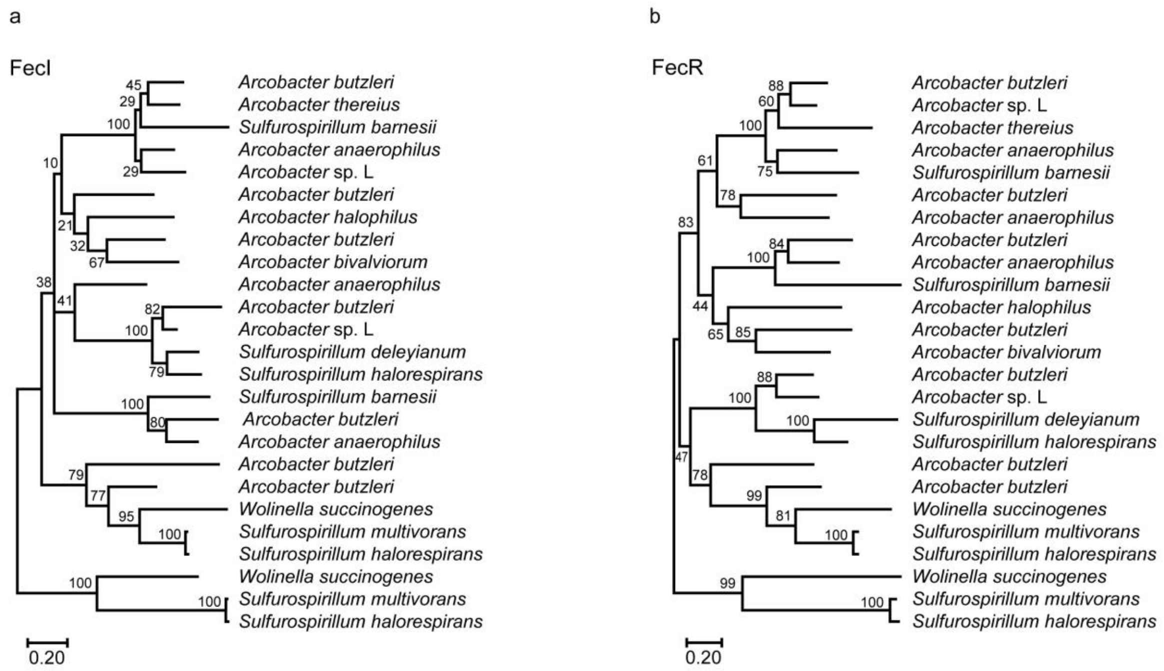


**Extended Data Fig. 6.** Phylogenetic tree of FecI (a) and FecR (b) proteins in the *Campylobacterota* phylum.

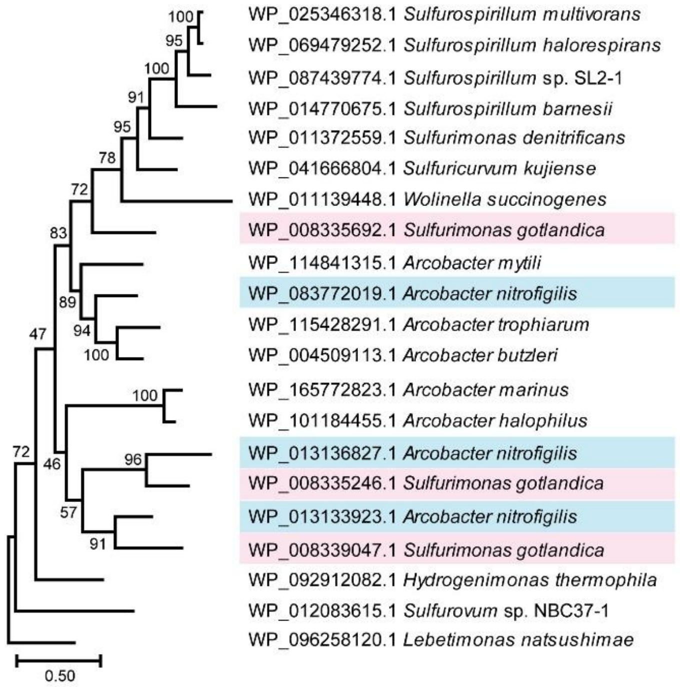


**Extended Data Fig. 7.** Phylogenetic tree of adenylate cyclase in the *Campylobacterota* phylum. Pink, 3 adenylate cyclase homologs in *Sulfurimonas gotlandica* GD1; Blue, 3 adenylate cyclase homologs in *Arcobacter nitrofigilis* DSM 7299.

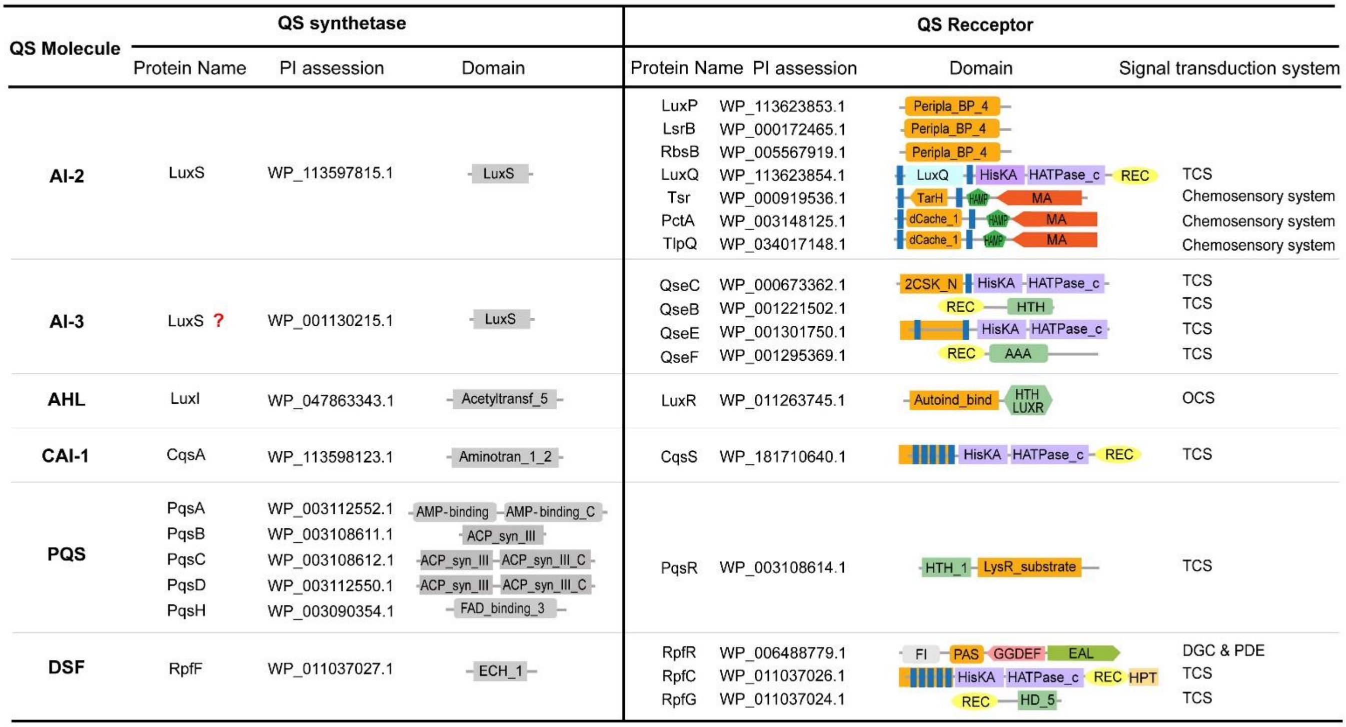

| QS Molecule | QS synthetase: Protein Name | PI assession | Domain | QS Recceptor: Protein Name | PI assession | Domain | Signal transduction system |
|---|---|---|---|---|---|---|---|
| **AI-2** | LuxS | WP_113597815.1 | LuxS | LuxP | WP_113623853.1 | Peripla_BP_4 | |
| | | | | LsrB | WP_000172465.1 | Peripla_BP_4 | |
| | | | | RbsB | WP_005567919.1 | Peripla_BP_4 | |
| | | | | LuxQ | WP_113623854.1 | LuxQ, HisKA, HATPase_c, REC | TCS |
| | | | | Tsr | WP_000919536.1 | TarH, HAMP, MA | Chemosensory system |
| | | | | PctA | WP_003148125.1 | dCache_1, HAMP, MA | Chemosensory system |
| | | | | TlpQ | WP_034017148.1 | dCache_1, HAMP, MA | Chemosensory system |
| **AI-3** | LuxS ? | WP_001130215.1 | LuxS | QseC | WP_000673362.1 | 2CSK_N, HisKA, HATPase_c | TCS |
| | | | | QseB | WP_001221502.1 | REC, HTH | TCS |
| | | | | QseE | WP_001301750.1 | HisKA, HATPase_c | TCS |
| | | | | QseF | WP_001295369.1 | REC, AAA | TCS |
| **AHL** | LuxI | WP_047863343.1 | Acetyltransf_5 | LuxR | WP_011263745.1 | Autoind_bind, HTH LUXR | OCS |
| **CAI-1** | CqsA | WP_113598123.1 | Aminotran_1_2 | CqsS | WP_181710640.1 | HisKA, HATPase_c, REC | TCS |
| **PQS** | PqsA | WP_003112552.1 | AMP-binding, AMP-binding_C | PqsR | WP_003108614.1 | HTH_1, LysR_substrate | TCS |
| | PqsB | WP_003108611.1 | ACP_syn_III | | | | |
| | PqsC | WP_003108612.1 | ACP_syn_III, ACP_syn_III_C | | | | |
| | PqsD | WP_003112550.1 | ACP_syn_III, ACP_syn_III_C | | | | |
| | PqsH | WP_003090354.1 | FAD_binding_3 | | | | |
| **DSF** | RpfF | WP_011037027.1 | ECH_1 | RpfR | WP_006488779.1 | FI, PAS, GGDEF, EAL | DGC & PDE |
| | | | | RpfC | WP_011037026.1 | HisKA, HATPase_c, REC, HPT | TCS |
| | | | | RpfG | WP_011037024.1 | REC, HD_5 | TCS |

**Extended Data Fig. 8.** Domain analyses of proteins in QS pathways. QS pathways are sorted by QS molecules; reported synthetases and receptors are listed in the table. The ligand binding domains for QS molecules in the receptors are highlighted in orange.
